# Enantiomer-Dependent Biological Activity of Cysteine-Coated Ceria Nanoparticles in Colorectal Cancer Cells

**DOI:** 10.64898/2026.04.27.721174

**Authors:** Emine Sümeyra Turali Emre, Ahmet Dinç, Sahra Esmkhani, Brendan Knittle, Nicole Sorensen, Ahsen Morva Yilmaz, Hülya Yazici, Hilal Yazici, Nicholas A. Kotov

## Abstract

Colorectal cancer (CRC) remains a major cause of cancer death, and advanced disease is still limited by resistance and systemic toxicity. We studied intrinsically active, biomimetic cerium oxide nanoparticles (CeNPs) functionalized with D- or L-cysteine (D-Cys@CeNPs and L-Cys@CeNPs) in three CRC cell lines (COLO-201, DLD-1, and LoVo) and healthy colon fibroblasts (CCD-18Co). We propose these materials act as enantioselective functional keys: cysteine stereochemistry shapes recognition at the nano–bio interface, while productive interactions allow the Ce³⁺-rich surface to drive localized redox exchange. We measured viability, ROS as a downstream phenotypic readout, Annexin V/PI-defined cell fate, and expression of the NF-κB regulatory genes TNFAIP3 (A20), IKBKG (NEMO), and NFKBIA (IκBα). Across the CRC panel, D-Cys@CeNPs caused earlier and stronger loss of viability, with the clearest effect in COLO-201, and shifted cells toward late apoptosis and necrosis. In contrast, L-Cys@CeNPs produced slower and more heterogeneous fate changes. Gene expression showed enantiomer-dependent differences in NF-κB feedback, consistent with differential pathway engagement. CCD-18Co fibroblasts were comparatively resistant to both enantiomers. Together, these findings link chiral CeNP surface design to redox-linked pathway regulation and support a materials-based route to selective anticancer activity.

INTRODUCTION

---

Colorectal cancer (CRC) is the third most frequently diagnosed malignancy and the second leading cause of cancer-related mortality worldwide [1–4]. Advances in screening, surgery, chemotherapy, targeted therapy, and immunotherapy have improved outcomes in selected patient groups. However, treatment options for advanced CRC remain limited by tumor heterogeneity, therapy resistance, and inadequate treatment selectivity. [4, 5]. These ongoing challenges underscore the need for therapeutic platforms capable of engaging tumor biology with high precision to minimize nonspecific damage to healthy tissues. [1, 6].

Nanomedicine [7, 8] specifically biomimetic nanoparticles (NPs), provides a robust framework to enhance therapeutic specificity, reduce systemic toxicity, and introduce novel medical functionalities [9–12]. Cerium oxide nanoparticles (CeNPs) are particularly notable for their inherent surface activity. The CeNP surface dynamically transitions between Ce³⁺ and Ce⁴⁺ oxidation states, enabling context-dependent redox exchange and catalytic functions. This switching capacity governs the catalytic behavior, electron-transfer efficiency, and biological reactivity of CeNPs [13–15]. Consequently, the balance of surface valence and the associated oxygen vacancy landscape dictate the material’s overall function [16].

In cancer research, nanoceria is frequently categorized simply as either a reactive oxygen species (ROS) scavenger or generator [17]. This binary view overlooks the strong influence of the local microenvironment, defect chemistry, and surface composition. A Ce³⁺-rich nanoparticle presents a highly defective and redox-responsive interface compared to a Ce⁴⁺-dominant particle. This physical distinction significantly alters how the particle interacts with surface biomolecules and intracellular signaling cascades [18–20]. Therefore, a Ce³⁺-rich chiral nanoceria system operates less as a basic source of oxidative stress and more as a localized, redox-active interface. It can predictably alter specific biomolecular interactions based on surface stereochemistry and cellular context [21].

Chirality introduces a critical second layer of selectivity. Biological systems are intrinsically chiral. Consequently, enantiomeric nanomaterials exhibit distinct interactions with proteins, membranes, and receptors, even when maintaining identical size, shape, and charge [22, 23]. Prior studies and reviews have shown that chiral nanoparticle surfaces can alter protein orientation, protein conformation, and receptor-associated cellular interactions, supporting the broader idea that chirality functions as a true biological design parameter rather than a decorative surface feature. For redox-active inorganic nanoparticles, this is especially important because stereochemistry can influence not only initial binding geometry but also the local interfacial environment in which electron transfer and signaling perturbation occur. For redox-active inorganic nanoparticles, stereochemistry influences initial binding geometry as well as the local interfacial environment where electron transfer occurs [24]. In a Ce³⁺-rich ceria system, the specific surface chirality derived from cysteine ligands directs how oxygen-vacancy-associated redox sites engage biomolecular targets. This process translates physicochemical asymmetry into distinct cellular responses [25–28].

In cell-based systems, total ROS measurements generally reflect an evolving cellular state as stress accumulates prior to fate decisions. Interpreting ROS as the sole initiating mechanism of cytotoxicity is therefore an incomplete model. A more robust approach connects the interface-defined redox state to specific pathway-level regulation and subsequent phenotypic outcomes [29–32]. In CRC, a primary pathway linking redox perturbations to cell survival or death is NF-κB signaling. This pathway integrates inflammatory, oxidative, and genotoxic signals to regulate apoptosis resistance and cellular proliferation [33]. Canonical NF-κB activity is controlled by specific regulatory nodes, including TNFAIP3 (A20), IKBKG (NEMO), and NFKBIA (IκBα), which manage pathway activation and termination [34]. Because these regulators operate within feedback loops, shifts in their expression provide a mechanistic readout of how redox-active interfaces influence survival signaling under cellular stress [4, 35].

Apoptosis is a highly regulated form of programmed cell death. t is the fundamental mechanism driving the efficacy of many anticancer agents, including chemotherapeutics (cisplatin, doxorubicin, 5-fluorouracil) and targeted therapies (TRAIL receptor agonists, Bcl-2 inhibitors) [36–39]. Conventional therapies often lack selectivity, leading to severe collateral damage in healthy tissues [40, 41]. Developing strategies that selectively induce apoptosis in cancer cells remains a major objective.

Nanoparticle chirality enables this targeted selectivity directly at the nano–bio interface, where biological surfaces govern contact formation, interaction persistence, and signal propagation through stereochemical recognition [42, 43]. In ceria systems, chirality regulates interaction probability while also controlling the resulting redox activity upon engagement. We propose a functional “enantioselective key” framework. In this model, cysteine stereochemistry dictates chiral recognition at the nano–bio interface by defining contact geometry. Upon productive binding, the Ce³⁺-rich surface initiates localized redox exchange. This key adapts to its environment: a single CeNP core presents different functional interfaces based on both the ligand stereochemistry and the accessible Ce³⁺/Ce⁴⁺ redox state. Together, these properties drive the enantiomer-dependent biasing of redox-linked signaling and NF-κB feedback loops. We capture this effect through the differential regulation of TNFAIP3, IKBKG, and NFKBIA, rather than relying on ROS generation as the primary initiating driver.

Here, we investigate D-cysteine- and L-cysteine-functionalized CeNPs (D-Cys@CeNPs and L-Cys@CeNPs) as intrinsically active biomimetic nanomaterials. We quantify their enantiomer-dependent effects on cytotoxicity, apoptotic phenotypes, and NF-κB regulatory gene expression (TNFAIP3, IKBKG, and NFKBIA). This work establishes an interface-centered framework that links chiral surface design and redox-active ceria chemistry directly to pathway-level regulation. The findings offer mechanistic insights into intrinsic nanoparticle function, guiding the design of highly selective targeted nanotherapeutics [33, 44].

## RESULTS and DISCUSSION

### 1. Characterization of D-Cys@CeNPs and L-Cys@CeNPs

We evaluated the physicochemical properties and verified the chiral surface functionalization of D-Cys@CeNPs and L-Cys@CeNPs using multiple analytical techniques. Transmission electron microscopy (TEM) demonstrated a uniform nanoparticle morphology. The measured particle sizes were 3.6 ± 0.5 nm for L-Cys@CeNPs and 3.9 ± 0.8 nm for D-Cys@CeNPs (**Figure 1A**). Dynamic light scattering (DLS) indicated average hydrodynamic diameters of 18 ± 5.08 nm and 15 ± 7.37 nm for L-Cys@CeNPs and D-Cys@CeNPs, respectively (**Figure 1B)**. Both formulations maintained low polydispersity indices (PDI ∼0.1–0.2), confirming narrow size distributions and robust colloidal stability in aqueous media. Zeta potential analysis showed negatively charged surfaces at physiological pH, yielding –28 mV for L-Cys@CeNPs and –25 mV for D-Cys@CeNPs (**Figure 1C**). We utilized circular dichroism (CD) spectroscopy to verify chiral functionalization. The spectra displayed distinct, mirror-image optical activity for D-Cys@CeNPs and L-Cys@CeNPs, confirming successful surface conjugation with enantiomerically pure cysteine ligands (**Figure 1D**).

**Figure 1:**
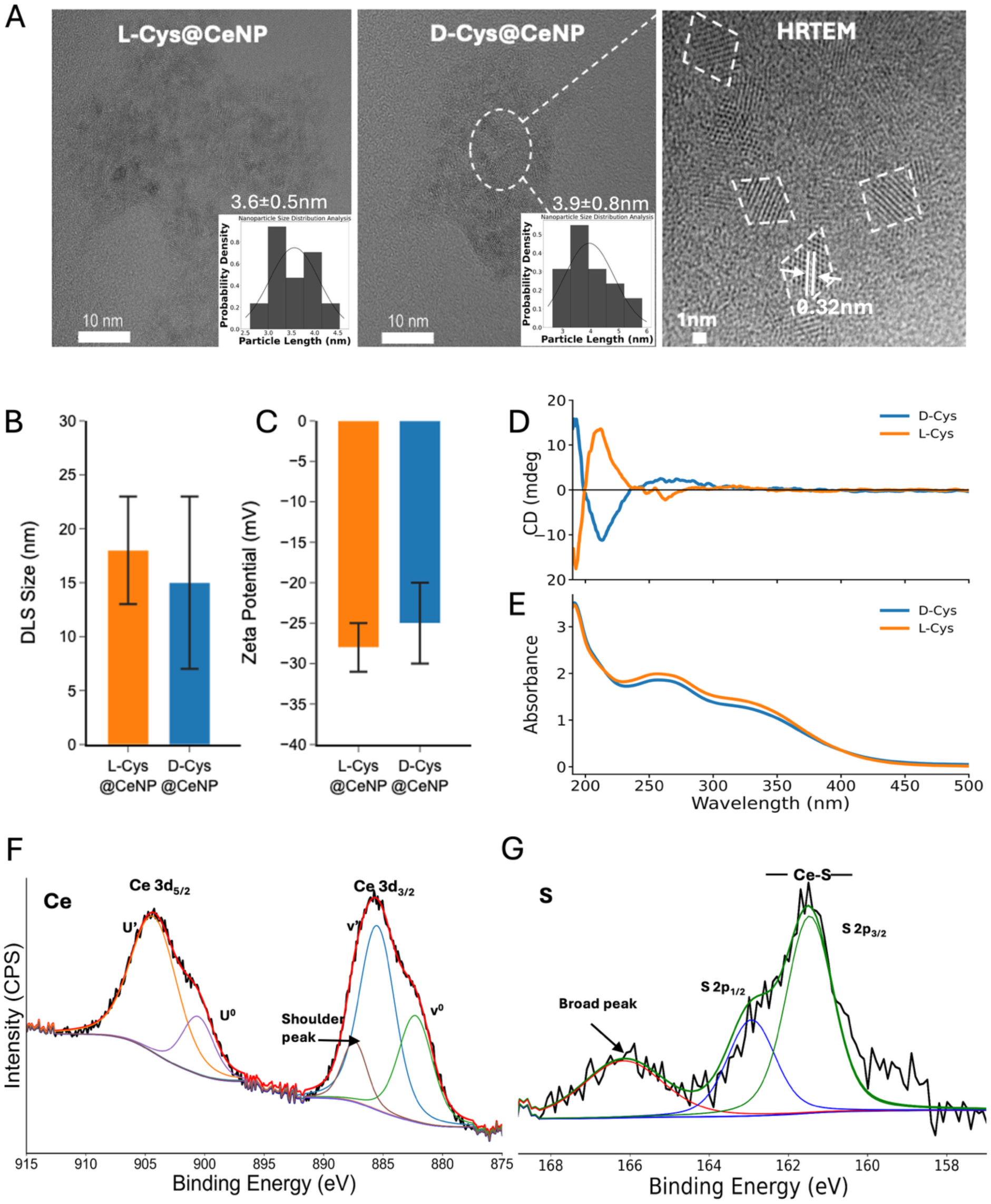
Characterization of NPs. (**A**) Transmission electron microscopy (TEM) images show the morphology of L-Cys@CeNPs (left) and D-Cys@CeNPs (right), with high-resolution magnification (inset, 1 nm scale) revealing crystalline features**. (B)** Dynamic light scattering (DLS) analysis shows narrow and comparable hydrodynamic diameter distributions for both enantiomers. **(C)** Zeta potential measurements indicate stable colloidal dispersion for both D- and L-functionalized CeNPs, with negative surface charge supporting electrostatic stabilization. **(D)** Circular dichroism (CD) and (E) UV-Vis absorbance spectra highlight the distinct chiroptical properties of D- and L-cysteine functionalized CeNPs, confirming successful surface chiral modification. **(F)** High-resolution Ce 3d XPS spectra for L-Cys@CeNPs indicate the presence of Ce³⁺ species, with peak deconvolution supporting a Ce₂O₃-dominant oxidation state. **(G)** High-resolution XPS spectra of the S 2p region confirm sulfur incorporation on L-Cys@CeNPs, consistent with thiol ligand binding.

High-resolution TEM (HRTEM) confirmed that the nanoparticles are highly crystalline structures with suggested truncated octahedral shape (**Figure 1A HRTEM**). This morphology stabilizes under the reducing synthesis conditions required for ultrasmall particle sizes, where the Ce³⁺ oxidation state dominates. Prior studies demonstrate that cerium oxide nanoparticles smaller than 5 nm favor bixbyite-type structures to accommodate oxygen deficiency and lattice rearrangement [45, 46]. The UV-Vis absorption profiles for both enantiomeric formulations are nearly identical, which aligns with expected optical behavior in non-polarized measurements **(Figure 1E).** The dominant spectral feature is a prominent absorption peak centered near 250–260 nm. This band corresponds to the allowed 4f¹→5d¹ electronic transition characteristic of Ce³⁺ ions [47]. In contrast, the O²⁻→Ce⁴⁺ charge transfer band, typically a broad and intense feature between 300 nm and 350 nm in Ce⁴⁺-rich systems, is suppressed [48]. The high intensity of the Ce³⁺ peak, combined with minimal absorption tailing in the 300-350 nm range, physically demonstrates a high concentration of Ce³⁺ relative to Ce⁴⁺.

X-ray photoelectron spectroscopy (XPS) was also employed to confirm the oxidation state of cerium. Deconvoluted high-resolution Ce 3d XPS spectrum of both D-Cys@CeNPs and L-Cys@CeNPs consisted exclusively Ce^3+^, displaying four well-resolved peaks located at ∼882.3 eV (Ce 3d₅_/_₂–v₀), ∼884.8 eV (Ce 3d₅_/_₂–v′), ∼900.8 eV (Ce 3d₃_/_₂–u₀), and ∼903.2 eV (Ce 3d₃_/_₂–u′), which are characteristic of Ce³⁺ species [49]. (**Figure 1F and Figure SI 1D**). The complete absence of Ce⁴⁺ satellite peaks near ∼920 eV definitively rules out the presence of oxidized CeO₂ phases Both D-Cys@CeNPs and L-Cys@CeNPs displayed a reproducible shoulder peak at ∼885.8–886.2 eV, immediately adjacent to the v′ component. This feature indicates Ce³⁺ ions existing in a modified surface coordination environment. We attribute this to ligand-induced electronic perturbations from thiolate (Ce–S) or carboxylate (Ce–OOC) bonding, supporting the model that surface Ce⁴⁺ is reduced to Ce³⁺ via charge transfer or oxygen vacancy formation coupled to Ce–S bonds [50]. The peak alignments at ∼884.8 eV and ∼903.2 eV match the expected Ce 3d₅_/_₂ and Ce 3d₃_/_₂ spin–orbit components for a bixbyite cerium(III) oxide lattice [46]. Measurements of the lattice fringes within individual crystalline domains revealed an interplanar d-spacing of 0.32 nm. This matches the expanded (111) lattice planes of a defect-fluorite cell, or the (222) planes of the C-type bixbyite Ce₂O₃ structure [51].

XPS data further confirmed the presence of cerium, sulfur, nitrogen, oxygen elements on the Elemental survey scans confirmed the presence of cerium, sulfur, nitrogen, and oxygen on the nanoparticle surfaces (SI 1A). High-resolution S 2p spectra for both L-Cys@CeNPs and D-Cys@CeNPs displayed a clearly resolved doublet at 161.9 eV (S 2p₃/₂) and 163.1 eV (S 2p₁/₂) **(Figure 1G and Figure SI 1F).** These binding energies indicate a metal-thiolate bond, confirming that cysteine anchors to the NP surface via its thiol group. A minor secondary component at ∼165.2 eV represents partially oxidized sulfur species [52], such as Ce–O–S linkages or sulfonates formed during synthesis. These structural features were consistent across both enantiomeric formulations. High-resolution C 1s spectra confirmed the structural integrity of the surface-bound cysteine (**Figure SI 1C**). The main peak at 284.8 eV was attributed to sp³-hybridized C–C and C–H bonds from the cysteine backbone (**Figure SI 1C-green**). A second peak at ∼286.0 eV represents C–N bonds from the amino group and potential C–O bonds from a hydroxylated surface. A third component at 287.9–288.8 eV aligns with the carboxyl carbon in either a protonated (–COOH) or coordinated (–COO⁻) state (**Figure SI 1C-red**). These features are typical for amino acids surface functionalization and confirm the presence of intact cysteine. While citrate was used as a stabilizing agent during synthesis, likely coordinating via weak carboxylate interactions, the stable chiral functionalization occurs predominantly through cysteine thiol binding. High-resolution O 1s spectra further verified a mixed oxygen environment spanning both the NP lattice and the surface ligands (**Figure SI 1B**). The primary peak at ∼528.3 eV was assigned to the Ce–O–Ce framework (**Figure SI 1B-red**). A secondary component at ∼529.0 eV aligns with surface hydroxyls (Ce–OH) and possible Ce–O–S bridging interactions [53]. A third peak at ∼532.8 eV represents oxygen within carboxylate groups (–COO⁻) (**Figure SI 1B-blue**). This data confirms a dual binding mechanism involving both thiol and carboxyl group coordination. The robust spectral fitting, characterized by low residual standard deviations (e.g., STD = 0.553 for S 2p, 0.938 for Ce 3d), strongly supports this structural model.

These collective findings confirm the successful synthesis and chiral surface functionalization of bixbyite-structured Ce₂O₃ nanoparticles. The resulting nanoparticles are stabilized by citrate and specifically conjugated with D- or L-cysteine ligands. They maintain a dominant Ce³⁺ oxidation state, show definitive evidence of Ce–S and Ce–OOC coordination, and display controlled surface chemical heterogeneity. Their uniform nanoscale dimensions, stable negative surface charge, and proven chemical stability make both D-Cys@CeNPs and L-Cys@CeNPs strong candidates for targeted biomedical applications.

### 2. Colon Cancer Cell-Lines

To comprehensively evaluate the therapeutic potential of D-Cys@CeNPs and L-Cys@CeNPs, we selected a cell line panel representing the progression of colorectal cancer (CRC) from healthy tissue to aggressive metastasis **(Figure 2)**. These models were curated to reflect a broad spectrum of genetic backgrounds, varying antioxidant capacities, and distinct regulatory profiles of the NF-B/TNFAIP3 (A20) axis. This selection facilitates a detailed analysis of how nanoparticle chirality and surface functionality interact with specific cellular architectures to dictate fate decisions.

**Figure 2:**
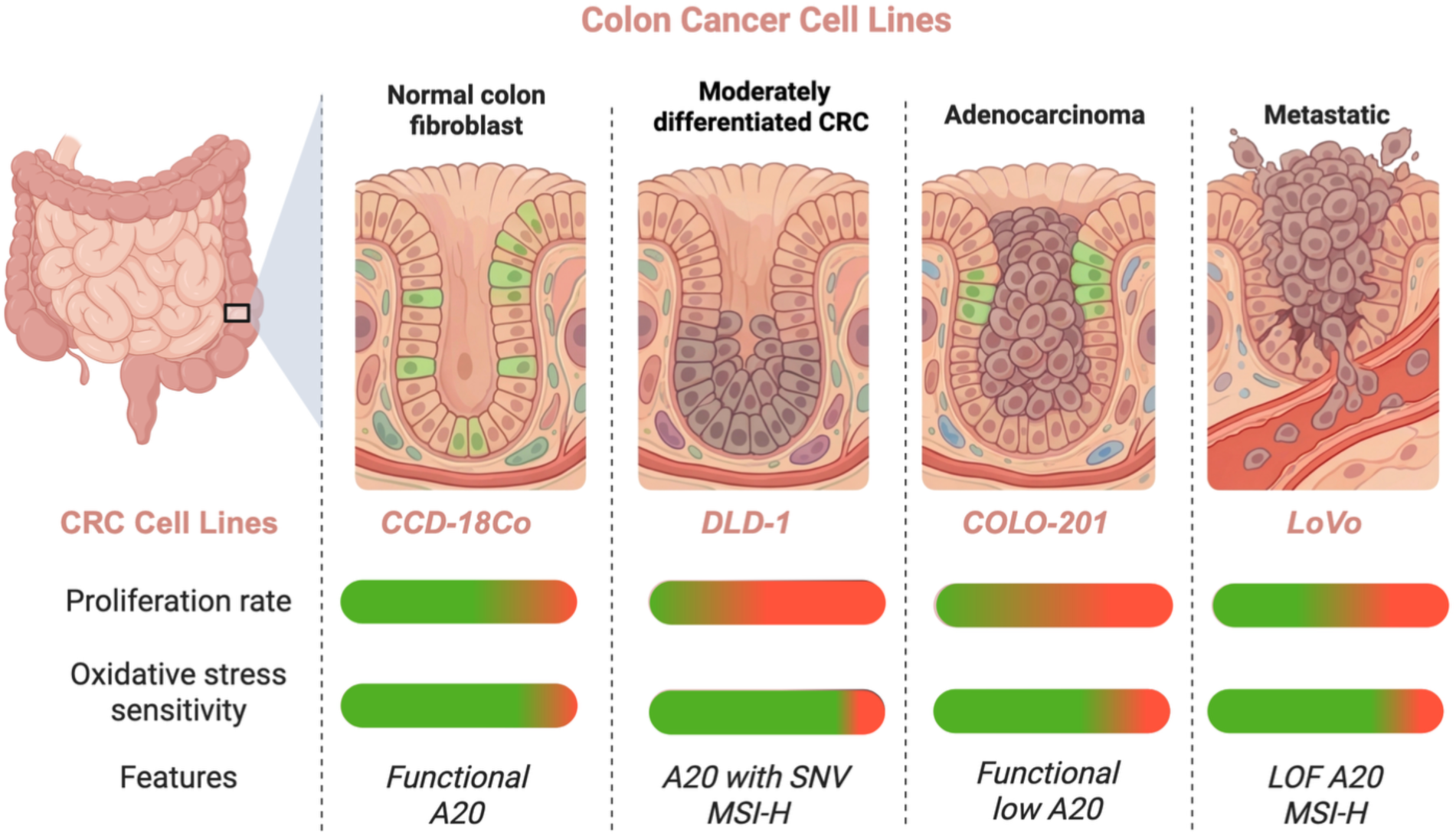
Phenotypic and genetic landscape of the colorectal cancer (CRC) panel. The selected models represent a clinical spectrum ranging from healthy colon tissue to aggressive metastasis. CCD-18Co serves as the non-malignant, healthy fibroblast control. COLO-201 is a highly proliferative adenocarcinoma characterized by high stress sensitivity and low functional A20 expression. DLD-1 is an MSI-H model with intact, albeit modified, A20 feedback (SNV:single-nucleotide variation). LoVo represents a metastatic MSI-H line harboring a Loss-of-Function (LOF) A20 mutation, maintaining resilience through a robust GSH-mediated antioxidant shield. (Gradient bars: Green = Low; Red = High). *Created with BioRender.com*.

#### CCD-18Co: Normal Colon fibroblast Control

The **CCD-18Co** line serves as the healthy control, representing non-malignant human colon fibroblasts. These cells exhibit low proliferation rates and maintain homeostatic redox regulation. Their signaling architecture and antioxidant pathways are well-balanced, providing a necessary reference for assessing therapeutic selectivity. This baseline ensures that any observed effects are grounded in cancer-specific dysregulations rather than non-specific toxicity, establishing a high safety threshold in the healthy stroma.

#### COLO-201: Adenocarcinoma Model

This cell line is a widely utilized adenocarcinoma model characterized by rapid proliferation and significant sensitivity to environmental stress[54]. At the molecular level, these cells are defined by a functional but low expression of A20, alongside limited antioxidant defenses, specifically evidenced by low intracellular glutathione (GSH) levels. Because the chemical buffers are minimal, the fate of the cell is heavily reliant on the stability of its remaining regulatory feedback loops. This makes COLO-201 a critical model for observing early-stage signaling intervention and high-affinity engagement at the regulatory node.

#### DLD-1: NF-κB Dysregulated, Microsatellite Stable CRC

**DLD-1** cells represent a moderately differentiated, Microsatellite Instability-High (MSI-H) CRC subtype. These cells harbor specific single-nucleotide variations (SNVs) in TNFAIP3 (p.A648S) and NFKBIA (p.P114Q), which subtly influence pathway activity without completely disabling the negative feedback loop. This intact regulatory machinery allows the cell to initially manage external stimuli and maintain signaling equilibrium. Consequently, **DLD-1** serves as a model for adaptive resilience, where functional regulatory “brakes” provide a temporary buffer against terminal signaling collapse.

#### LoVo: Metastatic CRC Model with High Antioxidant Capacity

The LoVo represents an aggressive metastatic **MSI-H** phenotype with a specific TNFAIP3 (A20) frameshift mutation, leading to a Loss-of-Function (LOF) state. This mutation potentially impairs A20 function, leading to enhanced, unchecked NF-B activation. Although this regulatory defect would typically render cells vulnerable, LoVo compensates through robust metabolic adaptation, characterized by significantly elevated GSH levels [55].

By incorporating these distinct CRC models, this study offers a comprehensive view of how chiral CeNPs interact with tumor cells across different biological backgrounds. This selection enables a rigorous assessment of how distinct biological backgrounds influence the transition between survival and apoptosis. By mapping the differential A20 status, ranging from low functional expression to total loss of function, we establish a comparative framework to evaluate the threshold for therapeutic engagement. This setup allows for a detailed analysis of whether the resulting cytotoxicity is a product of bulk oxidative damage or a more precise intervention within the cell’s internal signaling hubs.

### 3. Time- and Dose-Dependent Cytotoxicity and Therapeutic Selectivity of Chiral CeNPs

We evaluate the biological impact of D-Cys@CeNPs and L-Cys@CeNPs across CRC cell lines stated above (COLO-201, DLD-1, LoVo) and normal colon fibroblasts (CCD-18Co). The data presented in **Figures 3 and 4** demonstrate how nanoparticle chirality and treatment duration interact with specific cellular phenotypes to determine therapeutic outcomes.

**Figure 3.**
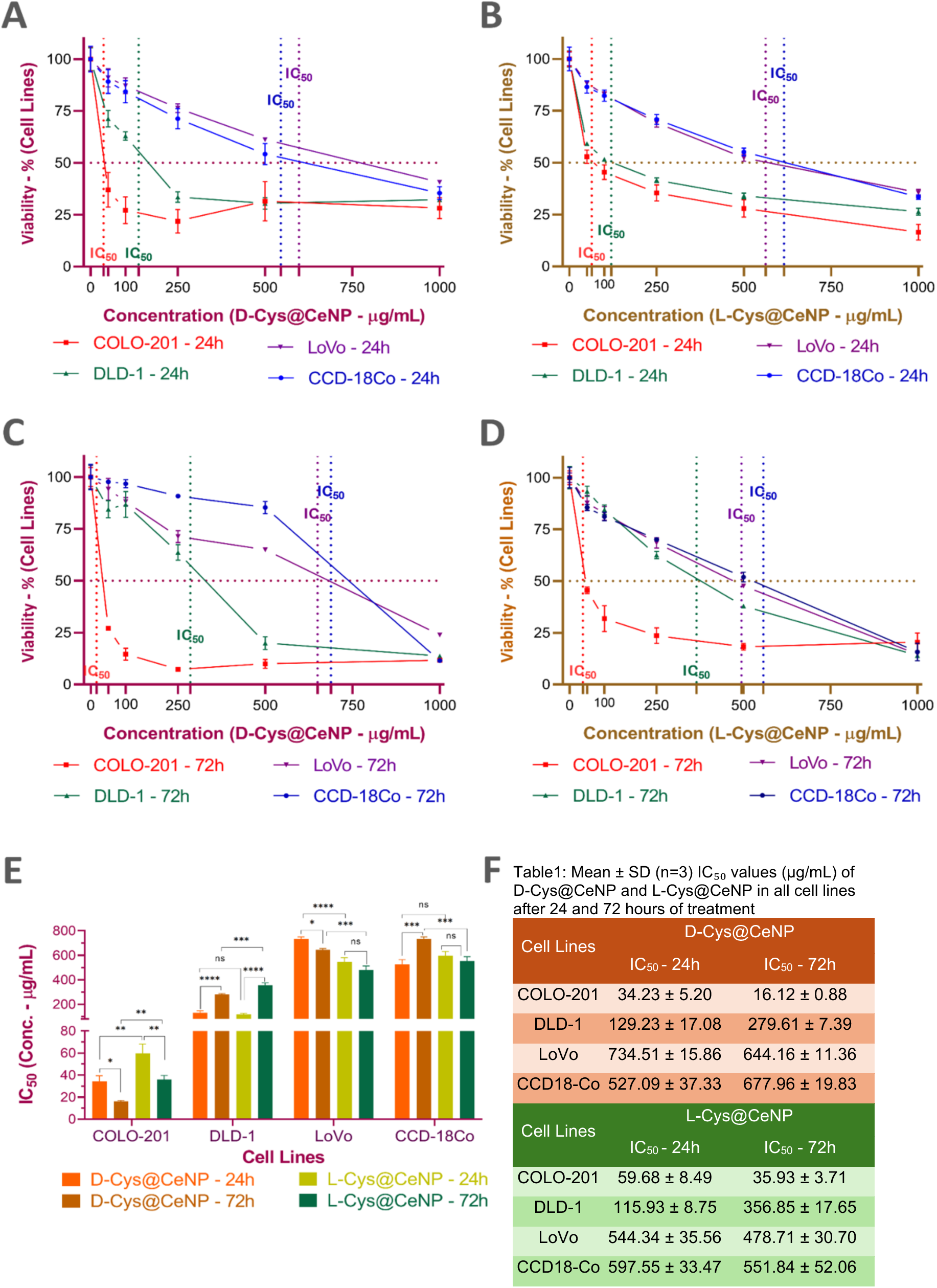
Comparative cytotoxic effects of D-Cys@CeNP and L-Cys@CeNP on different cell lines. **(A & B**): Cell viability of COLO-201, DLD-1, LoVo, and CCD-18Co cells treated with D-Cys@CeNP **(A)** and L-Cys@CeNP **(B)** for 24 h at increasing concentrations. **(C & D)** Cell viability of COLO-201, DLD-1, LoVo, and CCD-18Co cells treated with D-Cys@CeNP **(C)** and L-_34_C_0_ys@CeNP **(D)** for 72 h at increasing concentrations. **(E)** IC_50_ values for D-Cys@CeNP and L-Cys@CeNP treatments at 24 h and 72 h for all cell lines. **(F)** Summary table displaying the mean IC_50_ values (µg/mL) for each treatment condition across all cell lines. The results represent the mean ± SD of three independent biological replicates (n=3). The dotted vertical lines indicate the IC_50_ values, which are also displayed in the figure. Statistical significance was analyzed using one-way ANOVA followed by Tukey’s post-hoc test: (ns = non-significant, * p < 0.05, ** p < 0.01, *** p < 0.001, **** p < 0.0001).

**Figure 4:**
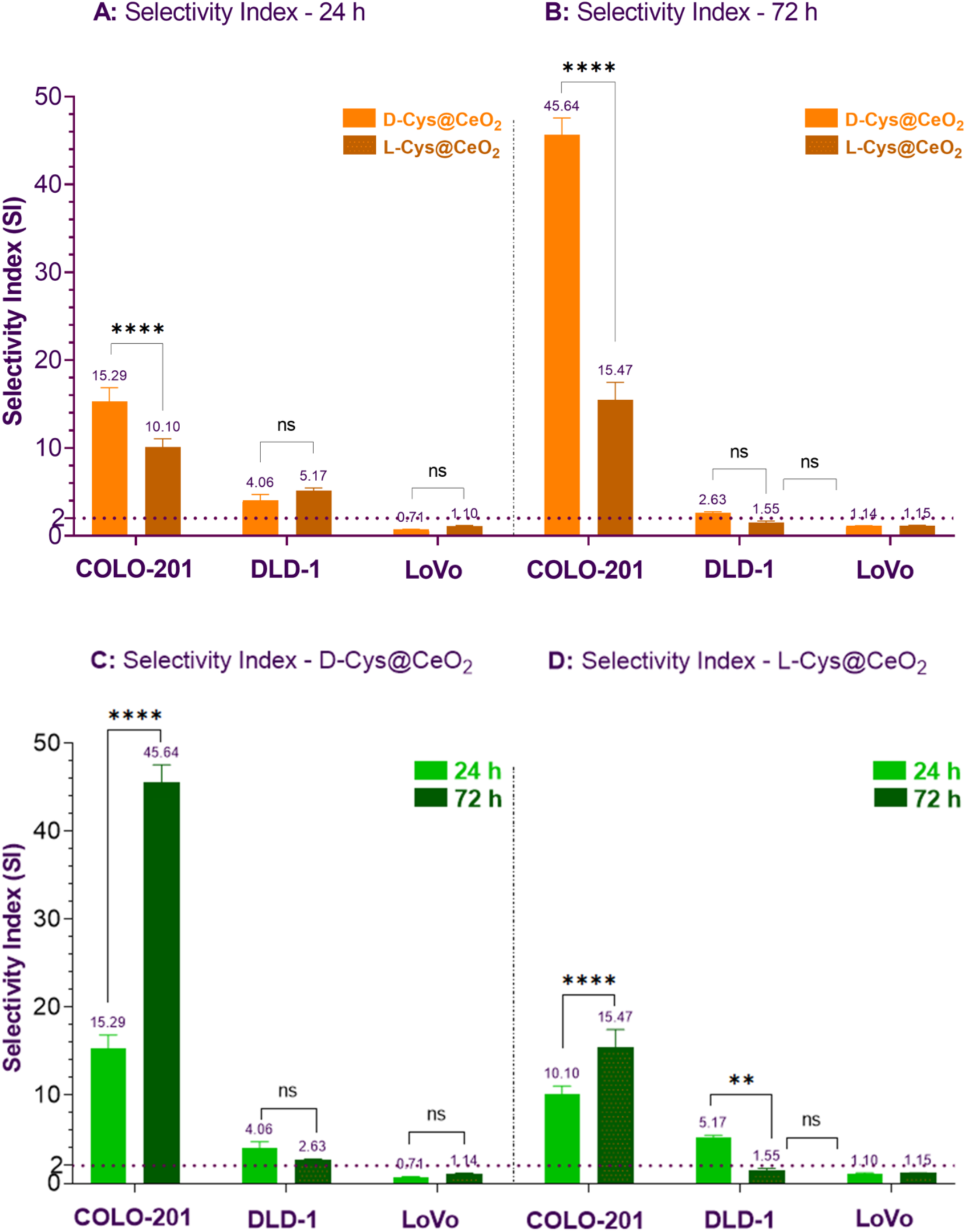
Selectivity Index analysis of D-Cys@CeNP and L-Cys@CeNP Across Cell Lines at 24 h and 72. **h.** The Selectivity Index is defined as the ratio of IC50 values in healthy control cells (CCD-18Co) to those in CRC cell lines (LoVo, DLD-1, and COLO-201). **(A)** Enantiomeric Comparison at 24 h: Comparison of SI values between D-Cys@CeNP (orange) and L-Cys@CeNP (brown) after 24 hours of exposure**. (B)** Enantiomeric Comparison at 72 h: Comparison of SI values between D-Cys@CeNP and L-Cys@CeNP after 72 hours of exposure. **(C)** Time-dependent analysis of D-Cys@CeNP selectivity across CRC cell lines. Over time NPs become more selective in COLO-201**. (D)** Time-dependent analysis of L-Cys@CeNP. The horizontal dashed line at SI=2.0 represents the baseline for selectivity. Data are presented as mean ± SD (n=3). Statistical significance is indicated by asterisks (, * p < 0.05, ** p < 0.01, *** p < 0.001, **** p < 0.0001)

**Figures 3** illustrates the concentration-dependent viability profiles (0 to 1000 g/mL) for both enantiomers. Each cell line is consistently represented by a color-coded trace: Pink (COLO-201), Green (DLD-1), Purple (LoVo), and Blue (CCD-18Co). Analysis of time- and enantiomer-dependent data for each cell is presented in **SI Section 2 (Figure SI 2-5)**.

#### Short-Term Response: 24-Hour Exposure (Figure 3A &B)

The D-enantiomer induced an immediate, aggressive cytotoxic response in the sensitive models **(Figure 3A- D-Cys@CeNPs)**. The COLO-201 (Pink) line showed a sharp decline at low concentrations, while the DLD-1 (Green) line reached ∼33% viability by 250 *μ*g/mL. In contrast, the CCD-18Co (Blue) healthy control-maintained viability above 90% at therapeutic doses, confirming an early-stage therapeutic window.

The response to the L-enantiomer was notably more gradual **(Figure 3B - L-Cys@CeNPs)**. While the COLO-201 (Pink) and DLD-1 (Green) lines still declined, the slopes were significantly less steep than those in **Figure 3A**. The LoVo (Purple) line exhibited strong initial resistance, remaining largely unaffected for 24 hours.

#### Long-Term Response: 72-Hour Exposure (Figure 3C&D)

Prolonged exposure of D-Cys@CeNP maximized the sensitivity of COLO-201 (Pink), shifting the curve further to low concentration (below 50 *μ*g/ml) **(Figure 3C)**. A critical observation was the “adaptive recovery” of the CCD-18Co (Blue) line: at 500 *μ*g/mL, the healthy fibroblasts showed higher viability than at 24 hours, suggesting they successfully neutralized the initial stress. Conversely, the DLD-1 (Green) line began to plateau, indicating that the surviving population activated resistance pathways to halt further decline. **Figure 3D (L-Cys@CeNPs):** This panel shows the cumulative impact of the L-enantiomer. Interestingly, the LoVo (Purple) line, which was highly resistant at 24 hours, eventually succumbed to prolonged exposure, with viability falling to 14% at the highest dose. This confirms that while LoVo is resistant to the rapid action of the D-enantiomer, it is vulnerable to the progressive accumulation of the L-form.

As explained in **section 2**, the specific genetic and phenotypic profile of each cell line dictates how nanoparticles interact with both surface and internal biomolecules **(Figure 2)**. This engagement likely explains the delayed cytotoxic response observed at 24 hours. At higher concentrations (≥500 µg/mL) and extended exposure (72 h), the significant rise in cytotoxicity suggests that sustained accumulation eventually overrides cellular defenses. This shift is likely driven by the exhaustion of antioxidant mechanisms (**Figure 3A, D**). Notably, D-Cys@CeNP-induced toxicity reached a plateau, while L-Cys@CeNPs showed a progressive increase. This divergence indicates enantiomer-specific differences in how these particles are processed or retained intracellularly. These results underscore how membrane trafficking and stereospecific interactions collaboratively shape the therapeutic response in metastatic cells [52].

#### Comparison of IC_50_ Values Across Cell Lines at 24 and 72 Hours

To further quantify the cytotoxic effects of D-Cys@CeNPs and L-Cys@CeNPs, we evaluated their half-maximal inhibitory concentration (IC_50_) values at 24 and 72 hours. This analysis provides a precise comparison of nanoparticle potency across different cell lines while highlighting temporal variations in cytotoxic responses. The results revealed distinct cytotoxicity profiles based on cell type and treatment duration, with lower IC_50_ values correlating with increased cytotoxicity in colorectal cancer cells, whereas healthy fibroblasts exhibited higher IC_50_ values, indicating greater resistance to nanoparticle-induced cytotoxicity **(Figure 3 E&F)**.

COLO-201 cells displayed the highest sensitivity to nanoparticles, with IC_50_ values significantly decreasing over time (D-Cys@CeNPs: 34.23 µg/mL to 16.12 µg/mL; L-Cys@CeNPs: 59.68 µg/mL to 35.93 µg/mL). This suggests progressive cytotoxicity due to rapid nanoparticle uptake and low antioxidant defenses. D-Cys@CeNPs exhibited stronger cytotoxic effects, likely due to higher oxidative stress induction. DLD-1 cells ranked second in sensitivity but developed resistance over time, with IC_50_ values increasing (D-Cys@CeNPs: 129.23 µg/mL to 279.61 µg/mL; L-Cys@CeNPs: 115.93 µg/mL to 356.85 µg/mL). This trend indicates an adaptive response, delaying cytotoxic effects until a critical threshold is reached, potentially due to high proliferative capacity. LoVo cells exhibited the highest resistance among cancer cell lines, with IC_50_ values decreasing slightly but remaining elevated (D-Cys@CeNPs: 734.51 µg/mL to 644.16 µg/mL; L-Cys@CeNPs: 544.34 µg/mL to 478.71 µg/mL). Their strong antioxidant capacity and metastatic nature may contribute to their delayed cytotoxic response. CCD-18Co healthy fibroblasts showed the highest IC_50_ values, indicating strong resistance (D-Cys@CeNPs: 527.09 µg/mL to 677.96 µg/mL; L-Cys@CeNPs: 597.55 µg/mL to 551.84 µg/mL). This suggests robust antioxidant defenses that mitigate nanoparticle-induced stress **(Figure 3 E&F).**

### 4. Comprehensive Analysis of Therapeutic Selectivity

To define the therapeutic window, we calculated the Selectivity Index (SI) as the ratio of the IC_50_ in healthy CCD-18Co cells to that in each cancer line **(Figure 4)**. D-Cys@CeNPs achieved a substantial selectivity advantage in COLO-201 cells, reaching an SI of 15.29 at 24 h. This level of selectivity significantly exceeds the conventional therapeutic threshold (SI>2), indicating that the D-enantiomer targets cancer-specific pathways rather than inducing indiscriminate cytotoxicity. The most striking observation occurs at 72 hours, where D-Cys@CeNPs exhibit a remarkable increase in selectivity, reaching an SI of 45.64 (p<0.0001) **(Figure 4C)**. This nearly 3-fold increase from the 24-hour mark suggests a potent, time-dependent targeting mechanism.

A comparison between the İtwo enantiomers reveals that D-Cys@CeNPs consistently outperform their L-counterparts in terms of selectivity **(Figure 4A & B)**. In COLO-201 at 72 hours, the SI for the D-enantiomer (45.64) is significantly higher than that of the L-enantiomer (15.47). This divergence demonstrates that the biological response is not merely a consequence of the redox activity of cerium oxide but is fundamentally driven by stereospecific recognition. The D-enantiomer likely achieves a superior geometric “fit” within the intracellular signaling machinery, enabling more effective pathway modulation at lower concentrations compared to the L-enantiomer.

In contrast, DLD-1 and LoVo models exhibit significantly lower SI values, often hovering near or below the selectivity threshold **(Figure 4A & B)**. The moderate SI values in DLD-1 reflect its adaptive resilience, likely due to its functional regulatory feedback. Meanwhile, the near-total lack of selectivity in LoVo (SI≈1.1) highlights the impact of metabolic shielding, where elevated glutathione levels likely passivate the nanoparticle surface and prevent specific target engagement.

### 5. Dose-Dependent ROS Dynamics and the Decoupling of Oxidative Stress from Cytotoxicity in response to D-Cys@CeNPs and L-Cys@CeNPs

To understand the mechanisms underlying the observed cytotoxic effects, we investigated whether oxidative stress serves as the primary driver or a secondary consequence of the therapeutic response. Cerium oxide nanoparticles are characterized by dual pro- and antioxidant properties, making a precise evaluation of reactive oxygen species (ROS) critical for interpreting their biological impact. By mapping ROS generation across the cell panel **(Figure 5, SI 6 & 7)** against respective IC_50_ thresholds, we decoupled bulk oxidative damage from targeted signaling intervention.

**Figure 5.**
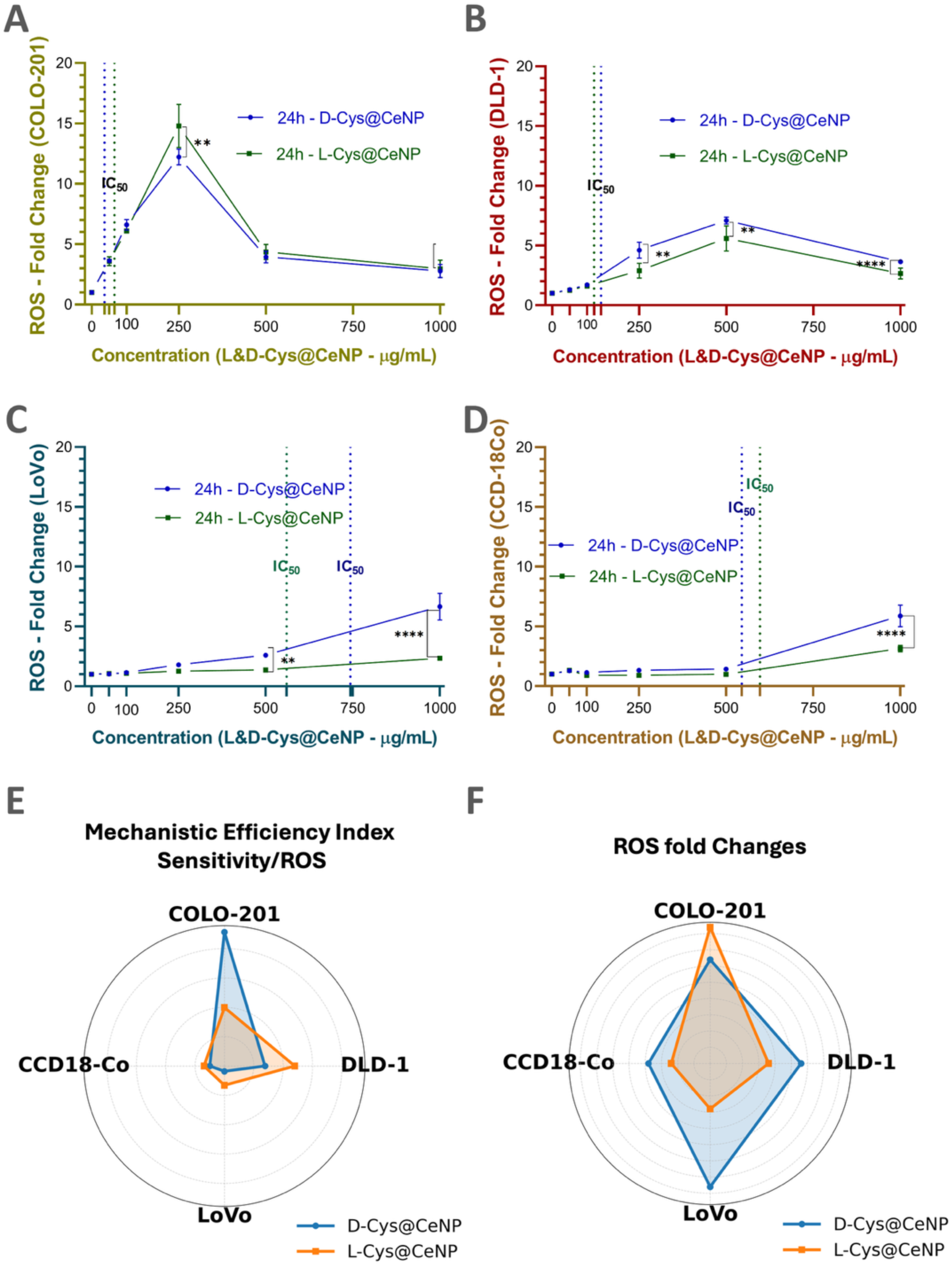
Characterization of ROS Dynamics and the Decoupling of Oxidative Stress from Cytotoxicity. **(A)** COLO-201 cells treated with D-Cys@CeNP and L-Cys@CeNP for 24 h; ROS fold change comparison. **(B)** DLD-1 cells treated with D-Cys@CeNP and L-Cys@CeNP for 24 h; ROS fold change comparison. **(C)** LoVo cells treated with D-Cys@CeNP and L-Cys@CeNP for 24 h; ROS fold change comparison. **(D)** CCD-18Co cells treated with D-Cys@CeNP and L-Cys@CeNP for 24 h; ROS fold change comparison. **(E)** Mechanistic Efficiency Index values for D-Cys@CeNP (orange) and L-Cys@CeNP (blue) across four cell lines: COLO-201, DLD-1, LoVo, and CCD-18Co**. (F)** ROS fold change at IC₅₀ concentrations in colorectal cancer and control cell lines treated with D-Cys@CeNP (red) and L-Cys@CeNP (blue) at their respective IC₅₀ concentrations.The vertical dashed lines indicate the IC_50_ values on the X-axis. * p < 0.05, ** p < 0.01, *** p < 0.001, **** p < 0.0001).

A fundamental observation across all evaluated cell lines is the absence of significant ROS accumulation prior to or at the IC_50_ thresholds **(Figure 5A–D).** If these nanoparticles-initiated cytotoxicity primarily through direct ROS generation, massive oxidative stress would necessarily precede or coincide with the IC_50_. Instead, substantial cell death occurs at relatively low concentrations (∼50–150 µg/mL) while ROS levels remain near baseline. ROS levels only surge at concentrations far exceeding the IC_50_, peaking at 250 µg/mL for COLO-201 and 500 µg/mL for DLD-1. This universal delay indicates that bulk oxidative damage is not the initiating mechanism. Early-stage cytotoxicity is instead attributed to the stereospecific engagement of the nanoparticles at the nano-bio interface, whereas subsequent ROS spikes represent a secondary consequence of terminal metabolic collapse.

The Mechanistic Efficiency Index **(Figure 5E)**, calculated as the ratio of cellular sensitivity (1/IC_50_) to total ROS generation, reveals a profound chiral divergence in the COLO-201 line. D-Cys@CeNPs exhibit an asymmetrical spike, achieving an efficiency value approximately 2.28 times higher than its L-counterpart. This demonstrates that the D-enantiomer eliminates COLO-201 cells with far greater precision per unit of ROS produced. This indicates a highly specific, non-redox-driven intervention at the signaling level, in which the nanoparticle serves as a functional key within a vulnerable biological environment.

The ROS fold-change profile **(Figure 5F)** contradicts the widely accepted hypothesis of death due to oxidative stress. In the sensitive COLO-201 model, L-Cys@CeNPs induce a higher absolute ROS response than D-Cys@CeNPs. If ROS were the primary executioner, the L-enantiomer would be the more potent formulation. Instead, the data clearly show that the D-enantiomer is significantly more lethal despite inducing less total oxidative stress. This proves that the high lethality of D-Cys@CeNPs in COLO-201 is independent of bulk ROS accumulation. The measured ROS is a secondary byproduct of the rapid collapse and mitochondrial failure initiated by the chiral NP intervention. Conversely, the resistant LoVo and healthy CCD-18Co cells exhibit delayed and lower ROS accumulation overall **(Figure 5C, D)**. In these lines, D-Cys@CeNPs induce higher absolute ROS than L-Cys@CeNPs, yet the Mechanistic Efficiency Index remains minimal. This confirms that while ROS can be generated in these environments, cells possess sufficient antioxidant shielding to resist ROS-mediated death.

It is important to note that the generation of ROS follows a dose-dependent, biphasic trajectory **(Figure 5 A&B)**. Sensitive models show a pronounced surge immediately following the IC_50_, followed by a sharp decline at 1000 µg/mL. At these massive excesses, the intrinsic ROS-scavenging capacity of the nanoceria dominates the microenvironment, neutralizing the oxidative stress generated by the dying population. This late-stage buffering protects the remaining viable fraction, explaining the slight recovery in cell viability observed in **Figure 3**.

### 6. Apoptosis Induction by D-Cys@CeNPs and L-Cys@CeNPs in Colon Cancer and Healthy Cell Lines

Following our cytotoxicity and ROS analyses, we wanted to determine whether the observed nanoparticle-induced stress responses ultimately lead to programmed cell death. We evaluated the extent of apoptosis induction by D-Cys@CeNPs and L-Cys@CeNPs in COLO-201, DLD-1, LoVo, and CCD-18Co cells using flow cytometry **(Figure 6)**. Each cell line COLO-201 **(Figure 6A)**, DLD-1 **(Figure SI 9)**, LoVo **(Figure SI 10)**, and CCD-18Co **(Figure SI 11)** was analyzed for apoptosis induction using Annexin V-PB/PI staining. Annexin V/PI flow cytometry categorized cellular states into: Viable (Q4: Annexin V−/PI−), Early Apoptotic (Q1: Annexin V+/PI−), Late Apoptotic (Q2: Annexin V+/PI+), and Necrotic (Q3: Annexin V−/PI+) population **(Figure 6A)**. This methodology distinguishes programmed cell death from unorganized membrane rupture by evaluating phosphatidylserine externalization and plasma membrane permeability. The scope of apoptosis induction by each nanoparticle type, along with variations in apoptotic responses across different cell lines, is detailed in the following sections. To characterize the dose-dependent transition in cell death mechanisms, cells were treated with nanoparticle concentrations spanning below, at, and above the established IC_50_ values. This gradient approach enables the differentiation between early signaling triggers and late-stage metabolic collapse, ensuring that observed enantioselective effects remain consistent across varying magnitudes of cellular stress. Furthermore, it confirms that the therapeutic response scales proportionally with nanoparticle accumulation rather than resulting from stochastic or localized events. **Figure 6A** displays representative flow cytometry data for COLO-201 highlighting the transition from viability to terminal death, while **SI 8-11** provides the statistical averages derived from five independent biological replicates for each cell line. This quadrant analysis facilitates a precise characterization of population shifts, confirming that the therapeutic response corresponds to the selective induction of programmed death pathways and subsequent secondary necrosis. The NPs’ apoptosis induction, along with variations in apoptotic responses across different cell lines, is detailed in the following sections.

**Figure 6:**
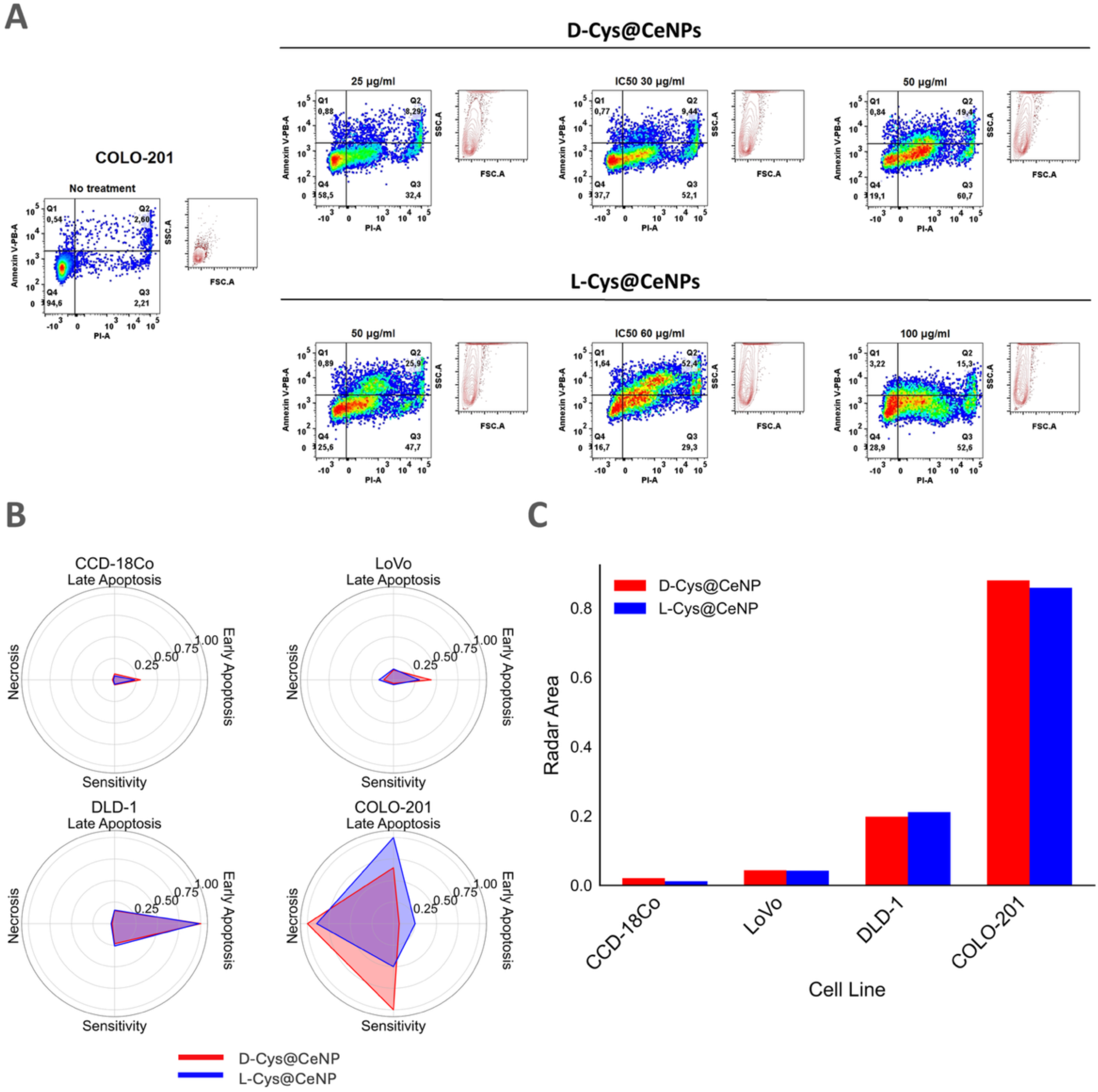
Flow cytometry-based apoptosis analysis. **(A)** Representative flow cytometry plots for each treatment condition in COLO-201 cells. Quadrant analysis distinguishes healthy (Q4; Annexin V-/PI-), early apoptotic (Q1; Annexin V+/PI-), late apoptotic (Q2; Annexin V+/PI+), and necrotic (Q3; Annexin V-/PI+) populations **(B-C)** Comparative analysis of apoptotic and cytotoxic responses induced by D-Cys@CeNP and L-Cys@CeNP in colorectal cancer (CRC) and healthy colon fibroblast cell lines at IC_50_ concentrations for 24 hours. **(B)** Cell line-specific radar plots comparing D-Cys@CeNP (red) and L-Cys@CeNP (blue) responses, highlighting differences in apoptotic phases and cytotoxicity profiles between the two nanoparticle enantiomers. **(C)** Bar plot comparing radar plot areas to represent the overall biological response magnitude for each nanoparticle in each cell line in B.

#### COLO-201 Cells

Treatment with D-Cys@CeNPs resulted in a dose-dependent reduction in cellular viability, characterized by a significant shift toward late apoptosis and necrosis (Figure 6A, SI 8). The quantitative population distribution (Table SI1) confirms the superior potency of the D-enantiomer and provides high-resolution insight into the kinetics of the death mechanism. At its IC_50_ (30 μg/mL), D-Cys@CeNPs reduced the healthy population to 44.44±4.58%, with the majority of cells transitioning to necrosis (37.68±10.04%) and late apoptosis (16.82±8.54%). In contrast, L-Cys@CeNPs required double the concentration (60 μg/mL) to achieve a comparable threshold (36.60±12.52% healthy).

The death profile for both enantiomers is defined by a rapid transition to terminal stages, as evidenced by the negligible early apoptotic populations (Q1), which remained below 2% for D-Cys@CeNPs at the IC50. Even at sub-IC50 concentrations (25 μg/mL), necrosis was already prominent (25.44±5.98%). This acute trajectory supports the “Functional Key” model: the signaling collapse initiated by stereospecific target engagement is likely too rapid to permit a prolonged early apoptotic phase. This interference appears sufficiently intensive to induce immediate mitochondrial dysfunction and ATP depletion, effectively “short-circuiting” the conventional apoptotic cascade [61]. Such metabolic failure directly correlates with the secondary necrosis and metabolic collapse observed in the prior ROS analysis.

Despite these shared trends, distinct chiral-dependent trajectories emerged. L-Cys@CeNPs predominantly induced late apoptosis, suggesting a more regulated and therapeutically relevant apoptotic pathway. Conversely, D-Cys@CeNPs exhibited elevated necrotic rates at 50 μg/mL, indicative of severe mitochondrial impairment [61]. These findings characterize L-Cys@CeNPs as potent inducers of controlled apoptosis, while D-Cys@CeNPs trigger an aggressive, mixed apoptotic-necrotic response driven by immediate signaling disruption [56]. Ultimately, the data confirms that while both enantiomers are potent therapeutic agents, D-Cys@CeNPs function as primary signaling disruptors, whereas L-Cys@CeNPs act as more conventional inducers of programmed cell death.

#### DLD-1 Cells

The population distribution in DLD-1 cells indicates a fundamentally different death trajectory compared to the acute signaling collapse observed in the sensitive COLO-201 model. At IC50 concentrations, the primary death signature is defined by a distinct shift toward Early Apoptosis (Q1), reaching 15.66 ± 7.87% for D-Cys@CeNPs and 15.31 ± 10.78% for L-Cys@CeNPs (Table SI2).

Unlike the rapid necrotic transition seen in COLO-201, the progression in DLD-1 remains largely confined to the early stages of regulated programmed cell death (Figure SI 9A). Notably, late apoptosis and necrosis markers remain minimal, generally staying below 5% even as concentrations increase to 250 μg/mL. This suggests that DLD-1 cells utilize an apoptosis resistance mechanism that prevents the “short-circuiting” of the mitochondrial pathway. This resilience is likely facilitated by known p53 mutations in the DLD-1 line, which inhibit caspase activation and conventional mitochondrial execution [1, 2, 63]. Furthermore, the activation of survival-related pathways, such as NF-κB signaling, likely delays apoptotic commitment by upregulating anti-apoptotic proteins [64].

The previously discussed ROS analysis corroborates this controlled response; while oxidative stress increases at IC50 doses, the absolute levels remain significantly lower than those in COLO-201 cells (Figure 5B). This indicates that DLD-1 cells regulate redox homeostasis more effectively to mitigate apoptosis induction. In terms of enantiomeric performance, DLD-1 exhibits a state of relative chiral indifference. While D-Cys@CeNPs predominantly trigger early apoptosis, L-Cys@CeNPs induce a comparable proportion of apoptotic cells, potentially due to minor differences in GSH metabolism interactions [65]. Overall, these findings suggest that while both nanoparticle formulations remain effective, the biological architecture of DLD-1 favors a slow, balanced apoptotic profile over the aggressive metabolic failure triggered in more vulnerable colorectal cancer lines

#### LoVo Cells

LoVo cells demonstrated pronounced apoptotic resistance, maintaining a high viable fraction (85–92%) throughout the nanoparticle concentration gradient **(Figure SI 10)**. Unlike the acute responses observed in COLO-201 and DLD-1, apoptotic markers remained remarkably low. Early apoptosis peaked at only 6.83% for D-Cys@CeNPs at the IC50, while late apoptosis stayed consistently below 5%. This resilience can largely be attributed to the MSI-H phenotype, which impairs DNA mismatch repair and disrupts conventional apoptotic signaling cascades [57, 58]. The FSC vs. SSC contour plots in **Figure SI 10A** reveal a significant morphological shift that provides a structural explanation for this sustained viability. While Annexin V/PI analysis confirms that the vast majority of cells remain in the healthy quadrant (Q4), the corresponding light-scattering profiles show a distinct, dose-dependent increase in Side Scatter (SSC). This increase in SSC correlates with enhanced intracellular granularity and structural complexity, indicating a high rate of nanoparticle internalization. LoVo cells appear to actively sequester Cys@CeNPs within endolysosomes or the cytoplasm without initiating programmed cell death. This confirms that these nanoparticles are biologically passivated, aligning with our Metabolic Shielding theory, where elevated GSH levels prevent the nanoparticles from engaging critical signaling “locks” [59].

This increased granularity is most prominent at the 1000 μg/mL concentration, where viability recovers to baseline levels (∼93–96% healthy). This suggests that high-density internalization is not a precursor to death but rather a mechanism of protection. During these massive excesses, the cerium oxide core acts as a high-capacity antioxidant reservoir, neutralizing endogenous reactive oxygen species in the cell environment [5, 6]. Consequently, the nanoparticles transition from potential “functional keys” to passive intracellular cargo. This decoupled relationship, where high internal granularity coexists with high viability, suggest the safety of the platform and suggests that these particles could be utilized as stable delivery vehicles in future studies.

Subpopulation analysis **(Figure SI 13)** further clarifies this heterogeneous response. Granulated cells exhibit a unique resilience, becoming more prevalent while maintaining high viability. In contrast, large-sized cells appear more susceptible to necrotic death, possibly due to a shift away from programmed pathways toward secondary necrosis and ATP depletion [72]. These findings suggest that LoVo cells utilize a combination of metabolic shielding and protective responses, such as autophagy, to mitigate oxidative stress and maintain homeostasis despite heavy nanoparticle loading [71].

#### CCD-18Co

CCD-18Co healthy colon fibroblasts **(Table SI4, Figure SI11)** establish the high biocompatibility and expansive therapeutic window of the Cys@CeNP platform. In stark contrast to the aggressive signaling collapse observed in sensitive cancer models, healthy cells exhibited near-total resilience to both enantiomers, even at supra-pharmacological concentrations.

Cellular viability remained exceptionally high throughout the experimental concentration gradient, with healthy populations consistently exceeding 90%. At their respective IC50 values (527 μg/mL for D-Cys and 598 μg/mL for L-Cys), the healthy fractions were measured at 92.58±1.54% and 94.70±3.03%, respectively. Even at the highest evaluated dose of 1000 μg/mL, viability remained stable at ∼90–94%, with negligible necrosis (∼1%).

The FSC vs. SSC contour plots **(Figure SI 11A)** reveal a dose-dependent increase in Side Scatter (SSC), particularly at 1000 μg/mL. Similar to the observations in LoVo cells, this morphological shift indicates a high rate of nanoparticle internalization and accumulation. However, because the Annexin V/PI profiles show no significant shift toward early or late apoptosis, it is evident that these internalized particles are biologically passivated. Healthy fibroblasts possess robust redox homeostasis and lack the vulnerable signaling hubs targeted by the nanoceria core, allowing them to host a heavy nanoparticle burden without compromising structural or functional integrity.

The SSC shift in CCD-18Co fibroblasts is notable but appears significantly less pronounced than the dramatic increase observed in the LoVo model. While there is a visible upward shift in internal complexity at 1000 μg/mL, the population remains more compact along the SSC axis compared to the extensively “stretched” profiles seen in LoVo. This difference suggests a more conservative rate of nanoparticle internalization or a more regulated accumulation process in healthy fibroblasts. Unlike cancer cells, which often exhibit upregulated endocytic pathways to meet high metabolic demands, healthy fibroblasts maintain more restrictive uptake kinetics.

These findings reinforce the safety of the Cys@CeNP platform through a double layer of protection. First, the platform is biologically passivated in healthy cells due to Metabolic Shielding and the absence of targeted signaling vulnerabilities. Second, the lower uptake kinetics in fibroblasts further minimizes the risk of physical stress or lysosomal overload. The nanoparticles are successfully delivered but remain therapeutically inactive confirming that the aggressive signaling disruption and subsequent metabolic failure observed in COLO-201 cells are exclusive to the cancer-specific “lock.” [60].

The observed increase in granular cell populations **(Figure SI 12A-C)** suggests that CCD-18Co cells may activate autophagy as a survival mechanism [61, 62]. These findings highlight those healthy fibroblasts exhibit strong resistance to CeNP-induced apoptosis due to antioxidant defense mechanisms, limited nanoparticle uptake, and other cytoprotective mechanisms.

#### Comparative Discussion on Different Mechanisms of Apoptosis According to Cell Lines

This study comprehensively compared apoptosis induction in COLO-201, DLD-1, LoVo, and CCD-18Co cells following CeNP treatment **(Figure 6)**. Radar plots constructed using values at the after 24 hours and integrated with a sensitivity axis, these geometric fingerprints reveal a clear hierarchy of death kinetics **(Figure 6B)**. While the resistant and healthy lines; DLD-1, LoVo, and CCD-18Co; remain primarily confined to the early apoptotic direction at their respective doses, COLO-201 is driven toward terminal stages. In this vulnerable line, the radar charts illustrate a stark chiral divergence: D-Cys@CeNPs aggressively favor necrosis, while L-Cys@CeNPs prioritize late apoptosis, signaling a total collapse of the biological defenses that the other lines still maintain.

The accompanying radar plot area graph is crucial for understanding the true magnitude of this selective engagement. By condensing multidimensional data potency and death mechanism into a single quantitative score, the area graph proves that the therapeutic impact is not just different in kind, but exponentially higher in degree for the targeted cancer model. The massive discrepancy in area between COLO-201 and the rest of the panel confirms that the **Cys@CeNP** platform is essentially silent in non-vulnerable environments.

COLO-201 cells exhibited high sensitivity to apoptosis, with L-Cys@CeNPs predominantly inducing late apoptosis, while D-Cys@CeNPs triggered necrosis due to severe mitochondrial impairment. In contrast, DLD-1 cells underwent early apoptosis but resisted late apoptosis and necrosis, likely due to p53 mutations, and effective ROS homeostasis maintenance. LoVo cells displayed strong apoptotic resistance, favoring autophagy and antioxidant mechanisms instead, suggesting that their MSI-H phenotype and high GSH levels contributed to their survival. Meanwhile, CCD-18Co cells demonstrated the highest resistance to apoptosis, with limited nanoparticle uptake and robust intrinsic antioxidant defenses. Overall, these findings highlight the necessity of tailoring CeNP-based therapies according to cancer type and apoptotic susceptibility, emphasizing the importance of cell-specific strategies to optimize therapeutic efficacy while minimizing off-target effects **(Table 2).**

**Table 2:** Comparative Summary of Apoptotic Profiles Induced by D-Cys@CeNPs and L-Cys@CeNPs.

| Cell Line | Apoptotic Sensitivity | Dominant Death Mode | IC<sub>50</sub> Response (Apoptosis) | Key Features |
| --- | --- | --- | --- | --- |
| COLO-201 | High | Late apoptosis and necrosis | L-Cys: 25.88% (60 µg/mL)<br>D-Cys: 18.55% (50 µg/mL) | Clathrin-mediated uptake, mitochondrial dysfunction. |
| DLD-1 | Moderate | Early apoptosis | D-Cys: 21.04% (250 µg/mL)<br>L-Cys: 18.96% (250 µg/mL) | p53 mutations, caspase pathway resistance, active NF-κB survival signaling. |
| LOVO | Low (Resistant) | Minimal apoptosis, some necrosis | D-Cys: 4.71% late apoptosis<br>L-Cys: <4% at all doses | MSI-H phenotype, elevated GSH, caveolar uptake, autophagy activation. |
| CCD-18CO | Very low (Selective sparing) | Minimal apoptosis | D-Cys: 2.81% late apoptosis (1000 µg/mL)<br>L-Cys: 1.23% late apoptosis (1000 µg/mL) | Low uptake, high antioxidant defenses, GSH preservation, survival through autophagy. |

These distinct apoptotic responses suggest an intricate balance between oxidative stress and survival pathways, which may be further influenced by NF-κB signaling. To elucidate how CeNPs modulate this critical regulatory network and its impact on apoptotic susceptibility, we next examined their effects on NF-κB activation, inhibition, and downstream gene expression in the cell lines.

### 7. Modulating NF-κB Signaling with Nanoparticles: Insights from Cancer Models

The regulatory effects of CeNPs on NF-κB signaling were investigated in COLO-201, DLD-1, LoVo, and CCD-18Co cell lines to evaluate their potential in modulating inflammatory and apoptotic pathways. Given NF-κB’s central role in tumor progression, therapy resistance, and immune regulation, the impact of CeNPs on its activation, inhibition, and downstream gene expression was assessed. The extent to which each nanoparticle type influenced NF-κB pathway components, alongside cell line-specific variations in signaling dynamics, is elaborated in the following sections. The canonical TNFR signaling pathway **(Figure SI14)** is a sophisticated molecular network that balances cellular survival and programmed death. The efficiency of this pathway is governed by three critical proteins, A20, NEMO, and IκBα which together form a functional regulatory axis that determines the duration, intensity, and ultimate outcome of NF-κB activation. These three proteins create a coordinated checks-and-balances system. facilitates the activation, gates the execution, and terminates the entire process. The functional synergy of this axis ensures cellular homeostasis. Any disruption, whether through genetic mutation, expression changes, or external modulation, alters the structural integrity of the pathway, shifting the balance from survival toward signaling collapse or persistent inflammatory activity.

#### COLO-201

The COLO-201 exhibits distinct responses to L-Cys@CeNP and D-Cys@CeNP treatments, demonstrating nanoparticle-specific impacts on NF-κB pathway regulation **(Figure7 red, SI 15)**. The low level of TNFAIP3 (A20) in COLO-201 is a critical determinant of its signaling dynamics, as this ubiquitin-editing enzyme plays a dual role as both a ubiquitin ligase and a deubiquitinase. Gene expression analysis revealed that L-Cys@CeNP treatment led to a moderate downregulation of TNFAIP3 (0.75-fold), suggesting incomplete suppression of NF-κB activation. This may indicate a threshold beyond which amplified A20 expression fails to fully counteract upstream activation signals. This phenomenon is supported by studies demonstrating that A20 overexpression does not significantly affect the NF-κB response, highlighting the limitations of A20’s regulatory capacity under persistent or strong activating conditions.[63] Conversely, D-Cys@CeNP significantly upregulated TNFAIP3 expression (2.60-fold), reflecting a robust activation of its feedback inhibitory function, which likely contributed to downstream NF-κB suppression [64]. IKBKG (NEMO), exhibited contrasting expression changes: it was upregulated with L-Cys@CeNP (2.12-fold) and downregulated with D-Cys@CeNP (0.75-fold). This suggests that L-Cys@CeNP activated NF-κB more strongly, potentially driving an inflammatory response, whereas D-Cys@CeNP suppressed this activation through enhanced A20-mediated feedback [65]. For NFKBIA (IκBα) both nanoparticles induced upregulation. However, L-Cys@CeNP treatment resulted in a stronger response (2.84-fold) compared to D-Cys@CeNP (1.20-fold), consistent with its heightened pathway activation. This elevated IκBα expression highlights the effectiveness of L-Cys@CeNP in triggering compensatory feedback mechanisms to maintain pathway homeostasis [66].

#### DLD-1

The DLD-1 colorectal cancer cell line demonstrated unique alterations in NF-κB signaling and apoptosis in response to L-Cys@CeNP and D-Cys@CeNP **(Figure 7, SI 15-purple)**. Mutations in *TNFAIP3* (p.A648S) and *NFKBIA* (p.P114Q) likely predispose the pathway to dysregulation [67]. These mutations impair the cell’s ability to terminate NF-κB activation and maintain feedback inhibition, contributing to the observed responses [68]. Gene expression analysis revealed that *TNFAIP3* (A20) was significantly downregulated in both treatments, with fold changes of 0.61 for L-Cys@CeNP and 0.33 for D-Cys@CeNP, suggesting a compromised ability to terminate NF-κB activation [69]. This effect was more pronounced with D-Cys@CeNP, indicating a greater disruption of regulatory mechanisms. Conversely, *IKBKG* (NEMO) was markedly upregulated, with fold changes of 3.18 for L-Cys@CeNP and 1.93 for D-Cys@CeNP, reflecting heightened NF-κB activation. NFKBIA (IκBα), a key feedback inhibitor, was also downregulated in both treatments (0.55 for L-Cys@CeNP and 0.64 for D-Cys@CeNP), further indicating impaired feedback inhibition that could exacerbate prolonged NF-κB activity [70].

**Figure 7:**
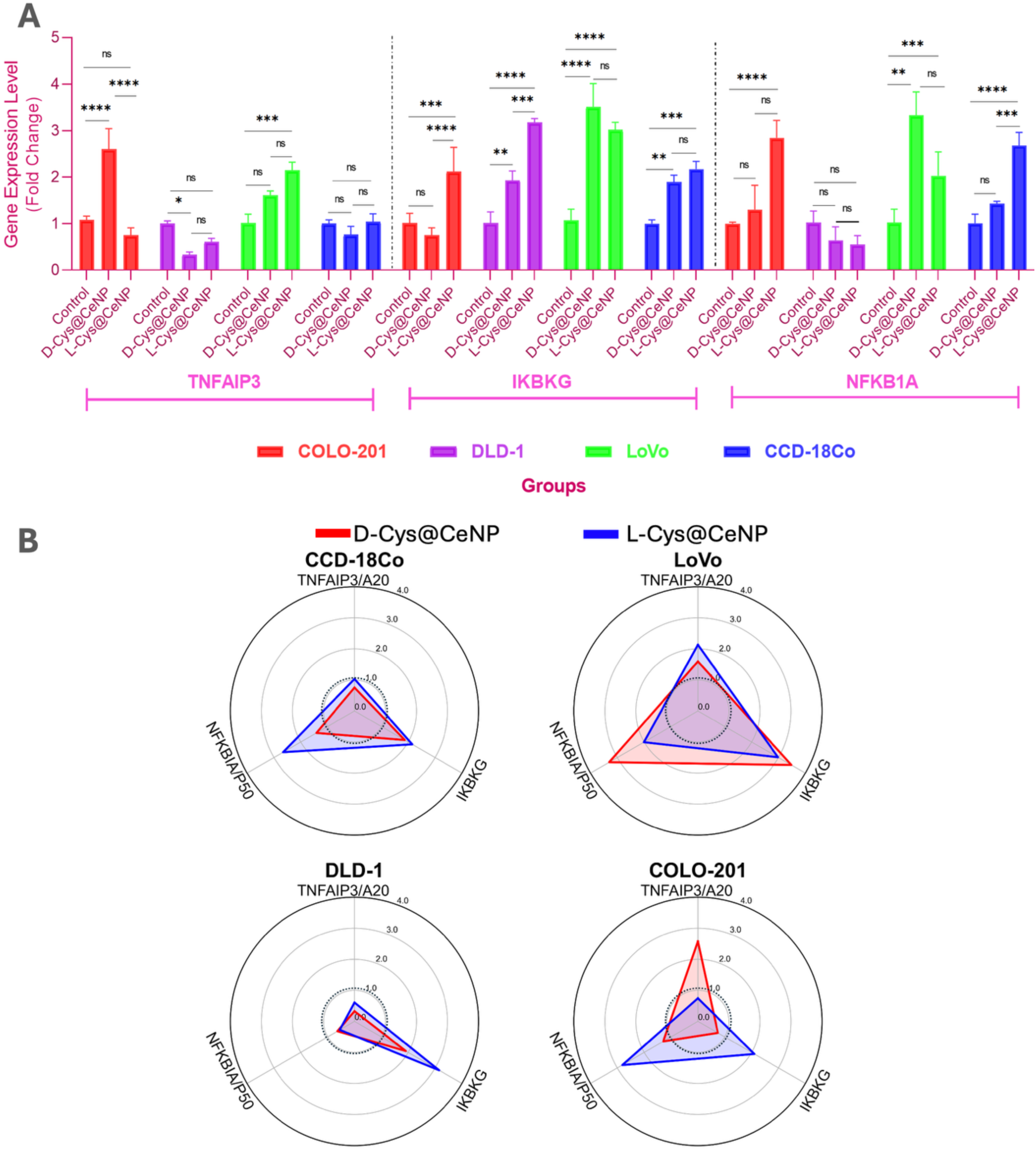
Gene expression analysis of TNFAIP3, IKBKG, and NFKBIA in colorectal cancer and normal fibroblast cells treated with D-Cys@CeNP and L-Cys@CeNP at IC_50_ concentrations for 24 hours. **(A)** Expression levels of TNFAIP3, IKBKG, and NFKBIA in all cell lines (COLO-201, DLD-1, LoVo, and CCD-18Co) after CeNP treatment. **(B)** Comparison of TNFAIP3, IKBKG, and NFKBIA expression within each cell lineand each nanoparticle treatment (D-Cys@CeNP vs. L-Cys@CeNP) across all cell lines. (ns = non-significant, * p < 0.05, ** p < 0.01, *** p < 0.001, **** p < 0.0001).

#### LoVo

The LoVo colorectal cancer cell line displayed a dynamic response to nanoparticle treatments, shaped by its complex genetic background, including a frameshift mutation in the TNFAIP3 (A20) gene **(Figure 7A-C-green)**. Both L-Cys@CeNP and D-Cys@CeNP treatments induced upregulation of TNFAIP3 expression, with a stronger response observed in L-Cys@CeNP treated cells (2.15-fold) compared to D-Cys@CeNP (1.61-fold). However, this frameshift mutation likely results in a truncated or functionally altered A20 protein, impairing its regulatory capacity [71]. Consequently, despite the upregulation of TNFAIP3 expression, the mutated protein may not fully suppress NF-κB activation. This partial loss of function could allow pro-survival signaling to persist, even under stress induced by nanoparticle treatments, ultimately enabling the cells to maintain their viability [72]. IKBKG (NEMO) was strongly upregulated, with D-Cys@CeNP eliciting a relativaly higher response (3.51-fold) compared to L-Cys@CeNP (3.01-fold). Similarly, NFKBIA (IκBα) showed greater upregulation with D-Cys@CeNP (3.33-fold) than L-Cys@CeNP (2.03-fold). The strong upregulation of IKBKG (NEMO) and NFKBIA (IκBα) following treatment with both L-Cys@CeNP and D-Cys@CeNP suggests robust activation of NF-κB signaling and the subsequent engagement of feedback inhibition mechanisms. The higher upregulation of IκBα with D-Cys@CeNP indicates a more pronounced feedback loop, which may help maintain NF-κB activity within a manageable range, preventing excessive apoptosis or necrosis [65]. Moving from cancer models to healthy fibroblasts reveals how genetic mutations influence NF-κB responses to Cys@CeNP.

#### CCD-18Co

In the healthy CCD-18Co colon fibroblast cell line, nanoparticle treatments elicited distinct effects on NF-κB signaling **(Figure 7, SI 15-blue)**. TNFAIP3 (A20) expression remained unchanged in both treatments, suggesting that the Cys@CeNP did not induce significant oxidative or inflammatory stress to activate this negative regulator. The stability of A20 implies that the NF-κB pathway activation was mild and within manageable levels, consistent with the cell line’s normal physiological state [73]. Conversely, IKBKG (NEMO) showed significant upregulation, with a fold change of 2.17 for L- Cys@CeNP and 1.89 for D- Cys@CeNP, indicating that both nanoparticles activated the canonical NF-κB pathway [4]. Similarly, NFKBIA (IκBα), a key feedback inhibitor, was markedly upregulated, with fold changes of 2.68 for L- Cys@CeNP and 1.43 for D- Cys@CeNP. This upregulation suggests an effective feedback mechanism preventing excessive NF-κB activity [74]. Overall, L-Cys@CeNP elicited stronger NF-κB activation compared to D-Cys@CeNP, as evidenced by higher NEMO and IκBα expression [65]. The results underscore the ability of CCD-18Co cells to maintain homeostasis under mild stress conditions through tightly regulated feedback mechanisms [75], with L-Cys@CeNP inducing a more pronounced but still controlled activation of NF-κB signaling.

#### Possible Mechanism of Cys@CeNP on NF-κB Pathway

Both L-Cys@CeNP and D-Cys@CeNP interact with the NF-κB pathway, eliciting distinct responses depending on the genetic background and mutation status of the cell lines. TNFAIP3 (A20), a key negative regulator of NF-κB, exhibited differential responses to nanoparticle treatment. In CCD-18Co cells, A20 expression remained stable, suggesting mild NF-κB activation that did not require significant feedback inhibition. In COLO-201 cells, where A20 is amplified, L-Cys@CeNP slightly downregulated A20, failing to fully suppress NF-κB activation, whereas D-Cys@CeNP robustly upregulated A20, effectively dampening NF-κB signaling. Despite a frameshift mutation in LoVo cells, A20 upregulation was still observed, indicating a partial loss of function that may sustain pro-survival signaling. In contrast, DLD-1 cells exhibited severe A20 downregulation with both treatments, reflecting a compromised negative regulatory capacity, leading to sustained NF-κB activation and heightened apoptosis. The regulators IKBKG (NEMO) and NFKBIA (IκBα) also showed cell line-specific responses. L-Cys@CeNP generally triggered stronger NEMO upregulation, promoting pathway activation and potential inflammatory signaling, whereas D-Cys@CeNP demonstrated a more controlled modulation by reducing NEMO expression in certain contexts, such as COLO-201. IκBα upregulation was more pronounced with L-Cys@CeNP, counteracting the heightened NF-κB activation, whereas D-Cys@CeNP appeared to rely more on A20-mediated suppression, particularly in COLO-201 cells. These findings suggest that CeNP-induced NF-κB modulation is highly dependent on the genetic landscape of each cell line, with D-Cys@CeNP exerting stronger feedback control through A20, while L-Cys@CeNP predominantly engages NEMO and IκBα-mediated regulatory mechanisms.

#### Link Between Gene Expression and Apoptosis Profiles

The apoptosis profiles of the cell lines align well with the gene expression data, highlighting the intricate role of NF-κB modulation in determining cell fate under nanoparticle-induced stress. The NF-κB pathway serves as a pro-survival transcription factor, capable of upregulating anti-apoptotic and pro-survival genes such as Bcl-2, Bcl-xL, and c-FLIP, which are critical for resisting apoptosis induced by moderate stress [76]. However, this pro-survival signaling is finely tuned by negative regulators such as IκBα (NFKBIA) and A20 (TNFAIP3), which ensure that NF-κB activation remains transient and context-dependent [73]. This checks-and-balances system is essential to prevent excessive or prolonged NF-κB activation, which can otherwise lead to cellular dysfunction.

#### COLO-201 Cells

The apoptosis data provide critical insights into how these gene expression changes translate to cell death outcomes. The significantly lower IC_50_ for D-Cys@CeNP (approximately half that of L-Cys) indicates its higher cytotoxicity. Apoptosis analysis revealed that COLO-201 cells treated with D-Cys@CeNP experienced markedly elevated levels of late apoptosis and necrosis, suggesting that the robust upregulation of A20 might shift the cell death mode from controlled apoptosis to necrosis under higher nanoparticle concentrations [77]. This aligns with the role of TNFAIP3 in inhibiting programmed cell death pathways while sensitizing cells to necrotic mechanisms under conditions of excessive stress or dysregulated signaling. In contrast, L-Cys@CeNP induced relatively lower apoptosis levels, consistent with its milder impact on TNFAIP3 expression and stronger NF-κB activation [78, 79].

#### DLD-1 Cells

The apoptosis results align with these gene expression findings. Both nanoparticles increased apoptosis levels, with D-Cys@CeNP exhibiting higher apoptosis percentages at all concentrations compared to L-Cys@CeNP. At concentrations near the IC_50_, the apoptosis rate for D-Cys@CeNP treated cells was significantly elevated, highlighting its stronger pro-apoptotic effect. This increased sensitivity to apoptosis in D-Cys@CeNP treated cells correlates with the severe downregulation of A20 and IκBα, which impairs the cell’s ability to counteract NF-κB-driven pro-survival signals [80, 81]. The slightly milder response to L-Cys@CeNP is consistent with its relatively less pronounced effects on these regulators.

#### LoVo Cells

The observed healthy apoptosis profile in LoVo cells, even at IC_50_ or higher concentrations of nanoparticle treatments, highlights the complex role of A20 in apoptosis regulation. While the frameshift mutation in the TNFAIP3 gene likely results in a truncated and functionally compromised A20 protein, residual activity or compensatory mechanisms may contribute to the observed response [82]. Additionally, certain studies suggest that A20 overexpression, even in its truncated form, might partially influence apoptosis regulation by interacting with components of the NF-κB pathway or other signaling networks [72]. Moreover, research has shown that A20 overexpression in glioblastoma contributes to resistance against TNF-related apoptosis-inducing ligand (TRAIL)-induced apoptosis [83], and A20 has been demonstrated to inhibit TNF-induced apoptosis in various cell lines [44]. However, the extent to which these findings apply to LoVo cells with a mutated A20 remains to be experimentally validated.

#### CCD-18Co Cells

Both nanoparticles maintain NF-κB activity within manageable limits, as indicated by stable A20 levels and significant IκBα upregulation. The apoptosis profile supports this, with the majority of cells remaining healthy, demonstrating that these cells effectively counterbalance stress induced by the Cys@CeNP.

While bar plots clearly show the magnitude of gene expression differences between D-Cys@CeNP and L-Cys@CeNP, radar plots reveal the overall shift in NF-κB pathway dynamics. Notably, COLO-201 cells exhibited strong TNFAIP3 induction upon D-Cys@CeNP treatment, indicating activation followed by feedback suppression, whereas L-Cys@CeNP induced greater IKBKG and NFKBIA expression in multiple lines, suggesting sustained NF-κB activation. Together, these data highlight enantiomer-specific modulation of inflammatory and apoptotic signaling.

Radar plots reveal cell-specific and enantiomer-dependent modulation of the NF-κB signaling pathway. Notably, **COLO-201 cells** showed strong induction of TNFAIP3/A20 in response to D-Cys@CeNP, while L-Cys@CeNP induced greater expression of IKBKG and NFKBIA in **CCD-18Co** and **DLD-1**, suggesting divergent regulatory effects on inflammatory and apoptotic signaling. Summarized gene expression changes, apoptosis profiles, and NF-κB pathway status following nanoparticle treatment are presented in **Table 3**..

**Table 3:**
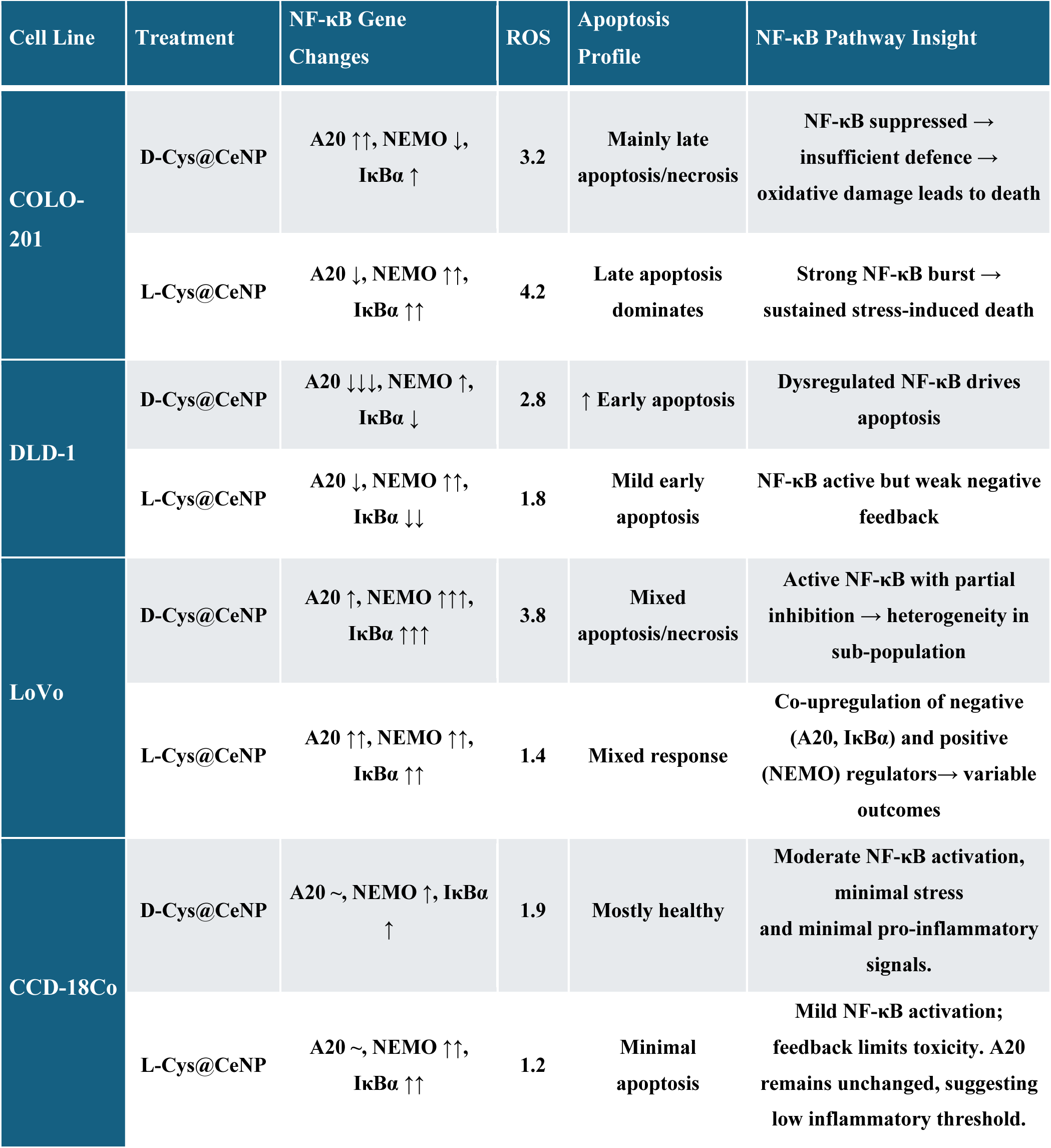
Summary of gene expression changes (TNAIP3, IKBKG, NFKBIA), apoptosis profiles, and NF-κB pathway status ollowing treatment with L-Cys@CeNPs and D-Cys@CeNPs in CCD-18co, DLD-1, COLO-201, and LoVo cell lines.

## CONCLUSION

The interaction between chiral nanoparticles and cellular interfaces appears to be a critical determinant of therapeutic outcomes in these colorectal cancer models. Our data support a “Functional Key” model, where D-Cys@CeNPs and L-Cys@CeNPs operate as stereospecific signaling modulators rather than simple pro-oxidants. The relationship between reactive oxygen species (ROS) and cytotoxicity is best understood through this framework, where ROS serves primarily as a downstream reporter of a committed apoptotic or necrotic collapse rather than its initiating cause. Ultimately, the therapeutic window is likely governed by the stereospecific fit at the nano-bio interface rather than the magnitude of the oxidative surge.

In vulnerable models such as COLO-201, the D-enantiomer appears to achieve a highly stereospecific geometric fit with the target complex. The canonical NF-κB pathway relies on a tightly regulated axis: NEMO serves as a requisite scaffolding component, IκBα acts as a repressor preventing unwarranted hyperactivation, and A20 operates as a negative feedback regulator constraining prolonged activation. Physical engagement by the nanoparticle positions its Ce³⁺-rich surface to exert localized redox modulation precisely at A20’s catalytic cleft. By potentially oxidizing the active-site cysteine within the triad [84], the nanoparticle effectively acts as a physical key “turning” the lock. Stripped of its deubiquitinase activity, A20 would fail to execute its negative feedback loop. Consequently, the signaling network abruptly collapses, short-circuiting pro-survival pathways and forcing the cell into terminal necrosis. This targeted signaling collapse kills with surgical precision, leaving a remarkably smaller “oxidative footprint” than its less-effective L-counterpart. The therapeutic efficacy, therefore, appears to be driven by the selective delivery of this localized oxidative payload directly to the catalytic cysteine of the A20 lock, functioning entirely independent of bulk oxidative stress.

Conversely, resistant cells lack an accessible biological lock, leading to distinct adaptive responses. Because DLD-1 possesses a functional, albeit mutated, A20 pool, these cells can initially buffer the nanoparticle’s interference. This buffering likely explains the observed shifts toward early apoptosis and results in a narrower therapeutic window compared to the A20-starved COLO-201. In more resilient models like LoVo, despite harboring a Loss-of-Function (LOF) A20 mutation, elevated GSH levels likely passivate the nanoparticle surface. This dense antioxidant buffer effectively blocks the specific “key-to-lock” interaction, explaining why even higher absolute ROS levels fail to trigger death in these environments.

The clear divergence between aggressive signaling disruption in vulnerable lines and passive internalization in resistant ones highlights a strategic opportunity. The recovery of viability and internal granularity observed at high doses in resistant lines and healthy CCD-18Co fibroblasts points to a biphasic role, where the nanoceria core may transition from a pro-apoptotic agent to a cytoprotective antioxidant reservoir. The ability of these cells to host a high nanoparticle burden without toxicity indicates that this platform could be utilized as a stable carrier for secondary molecular cargo.

Together, these findings establish that chiral nanoceria function as high-precision signaling modulators. Future studies integrating genomic and proteomic data with these geometric death profiles will be essential to fully map the enantioselective therapeutic window. By exploiting the unique redox properties and chirality of cerium oxide nanoparticles, we lay a foundation for more personalized treatment strategies. This approach may ensure that the potent redox capabilities of the Ce³⁺-rich surface are only unleashed within the targeted, cancer-specific lock. Refining these formulations for combination therapies could help overcome tumor resistance, further enhancing the specificity of nanoparticle-based interventions and offering a promising materials-based approach to colorectal cancer therapy.

## MATERIALS & METHODS

### 1. L-Cys@CeNPs and D-Cys@CeNPs Syntheses and Characterization

Chiral cerium oxide nanoparticles were synthesized using cerium nitrate and D- or L-Cysteine as surface ligands with water as the reaction medium. The reaction mixture contained final concentrations of 10 mM cerium nitrate, 10 mM D- or L-Cysteine, 10mM sodium citrate and 15 mM sodium borohydride. After two hours of stirring at room temperature, yellowish nanoparticle dispersions were obtained. Nanoparticles were purified by precipitation with isopropanol, followed by centrifugation at 4000 rpm for 15 minutes, washed with water, and lyophilized for storage. The nanoparticles were dissolved in MilliQ water or media for further analysis or cell experiments.

To assess the morphology and size of the nanoparticles, Transmission Electron Microscopy (TEM) imaging was performed using a Thermo Fisher Talos F200X G2 Scanning/Transmission Electron Microscope (S/TEM) with an accelerating voltage of 200 kV. For TEM measurements, samples were prepared by placing a diluted aqueous nanoparticle suspension onto carbon-coated copper grids and allowing them to dry at room temperature. Average particle size is calculated using ImageJ analysis.

To evaluate the chiral properties of the nanoparticles, Circular Dichroism (CD) spectroscopy was carried out at 25°C with a JASCO J-815 CD. Spectra were recorded in the 190–800 nm. CD spectra were obtained by a JASCO J-815.

Dynamic Light Scattering (DLS) was employed at 25 °C to measure the hydrodynamic diameter and polydispersity index (PDI) of the nanoparticles in aqueous dispersion. These measurements were used to assess the size distribution and stability of the nanoparticles. The PDI is calculated as the square of the standard deviation divided by the square of the mean diameter.

Zeta potential data were measured at 25°C using a Zetasizer Nano ZS instrument (Malvern Instruments Ltd, Malvern, Worcestershire, UK). These measurements provided information on the surface charge and stability of the nanoparticles in suspension.

To assess the surface chemistry of the nanoparticles, X-ray Photoelectron Spectroscopy (XPS) was performed using a Kratos Analytical AXIS Ultra spectrometer. Particle suspensions were drop-cast onto silica wafers for XPS analysis, with the silicon (Si) signal used for charge shift calibration. The XPS spectra were deconvoluted and fitted to standard spectra with a shift in the standard deviation (STD) lower than 1.2.

### 2. Cell Lines

To evaluate the cytotoxic and anticancer activities of D-Cys@CeNPs and L-Cys@CeNPs, the following colorectal cell lines were used: CCD-18Co (ATCC), a healthy colon fibroblast cell line, was maintained in Eagle’s Minimum Essential Medium (EMEM, Sigma-Aldrich, St. Louis, MO, USA, M4655). The colorectal cancer cell lines DLD-1 and COLO-201 (ATCC) were maintained in RPMI-1640 (Sigma-Aldrich, St. Louis, MO, USA, R8758), while LoVo (ATCC) was cultured in F-12K Medium (HyClone, Cytiva, USA, Cat# SH30526.0). Following cryopreservation, the vials were transferred to liquid nitrogen storage under standardized conditions to ensure long-term viability. Prior to experimental use, frozen cells were rapidly thawed at 37°C, gently resuspended in their respective culture media, and centrifuged to remove residual DMSO. The cells were then seeded at an appropriate density and allowed to recover for 24 hours before further experimentation to ensure optimal viability and physiological function. Morphological assessment and viability testing using trypan blue exclusion were performed to confirm cell integrity post-thawing. Mycoplasma contamination was routinely monitored in cell cultures using two distinct detection methods: the MycoAlert™ PLUS Mycoplasma Detection Kit (Lonza, Cat#: LT07-703) and the e-Myco™ Mycoplasma PCR Detection Kit (Intron Biotechnology, Cat#: 25236). These tests were applied every 4–5 passages to assess potential contamination and were also performed before cryopreserving cells in liquid nitrogen storage to prevent cross-contamination.

### 3. Cytotoxicity (WST-1) Assay

Cells were seeded at a density of 5,000 cells per well in 96-well plates with a final volume of 100 µL per well. Each experiment was performed in triplicate for all cell lines. The plates were incubated at 37°C with 5% CO_2_ for 24 hours. Subsequently, D-Cys@CeNPs and L-Cys@CeNPs were added to the wells at concentrations of 50, 100, 250, 500, and 1000 µg/mL in a final volume of 200 µL per well. After 24 and 72 hours of exposure, the culture medium was removed to evaluate the effects of the nanoparticles on cell viability. A 1:10 diluted WST-1 reagent (REF: 11644807001, Roche, USA) was added at 100 µL per well, followed by a 2-hours incubation at 37°C with 5% CO_2_. Absorbance was measured at 450 nm, with background absorbance at 650 nm subtracted, using a Cytation 5 instrument (BioTek, USA). Cell viability was calculated relative to untreated controls (set as 100% viable). IC_50_ values were obtained from three independent experiments for each cell line. Details on IC_50_ calculation are provided in the supplementary file.

#### 3.1 Selectivity Index (SI) Calculation

To assess the therapeutic selectivity of D-Cys@CeNP and L-Cys@CeNP formulations, the Selectivity Index (SI) was calculated by comparing their cytotoxicity in cancerous versus non-cancerous cells. The SI quantifies how selectively a nanoparticle targets cancer cells while sparing healthy cells, with higher values indicating greater selectivity.

The SI was determined using the following equation:

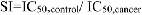

Where:

- IC₅₀ (CCD-18Co) is the half-maximal inhibitory concentration in the normal colon fibroblast cell line (CCD-18Co), and
- IC₅₀ (cancer) is the IC₅₀ value obtained from colorectal cancer cell lines (COLO-201, DLD-1, and LoVo), calculated independently for each nanoparticle formulation and time point (24 h and 72 h).

SI values were calculated for both D-Cys@CeNP and L-Cys@CeNP. An SI value greater than 2 was interpreted as an indication of preferential cytotoxicity toward cancer cells.

### 4. Reactive Oxygen Species (ROS) Assay

Cells were seeded at a density of 5,000 cells per well in 96-well plates with a final volume of 100 µL per well, and incubated for 24 hours at 37°C with 5% CO_2_. D-Cys@CeNPs and L-Cys@CeNPs were added at concentrations of 50, 100, 250, 500, and 1000 µg/mL, adjusting the final volume to 200 µL per well. After 24 hours of incubation, the culture medium was removed, and the cells were washed with 1**×** DPBS. A 10 µM solution of DCFDA (2′,7′-Dichlorodihydrofluorescein diacetate, Sigma Aldrich, D6883) was prepared in 1**×** DPBS and added to the wells at a volume of 100 µL per well. Plates were incubated at 37°C with 5% CO_2_ for 45 minutes. Fluorescence was measured at an excitation wavelength of 485 nm and an emission wavelength of 535 nm using a Cytation 5 instrument. ROS levels were expressed as fold change relative to the untreated control, normalized to the viability ratio at corresponding concentrations.

ROS levels were measured by DCFH-DA fluorescence at the IC₅₀ concentration of each nanoparticle after 3 hours of treatment, expressed as fold change relative to untreated controls. The resulting index (units: 1/ (µg mL⁻¹) integrates both oxidative stress induction and effective dosage, with higher values indicating greater ROS-driven cytotoxicity at lower nanoparticle concentrations.

#### 4.1 Calculation of Sensitivity and Mechanistic Efficiency Index (MEI)

To quantitatively assess the oxidative stress-mediated cytotoxic potential of chiral cerium oxide nanoparticles, a sensitivity coefficient and composite index were calculated for each cell line and treatment condition. The sensitivity coefficient was defined as the reciprocal of the IC₅₀ value:

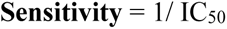

where IC₅₀ (µg/mL) represents the nanoparticle concentration required to reduce cell viability by 50%, as determined from dose–response curves (nonlinear regression, n=3).

To evaluate the cytotoxic efficacy of chiral CeNPs relative to their oxidative output, we calculated a Mechanistic Efficiency Index (MEI). This parameter isolates targeted lethal potency from non-specific oxidative initiation. Sensitivity was defined as the inverse of the half-maximal inhibitory concentration derived from 24h cytotoxicity assays.

The MEI was calculated using the following formula:

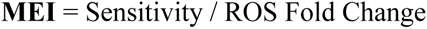

This index differentiates between ROS-mediated cell death (low MEI) and targeted, ROS-independent programmed cell death (high MEI).

### 5. Flow Cytometry-Based Apoptotic Assay

Cells were seeded in 6-well plates at a density of 100,000 cells per well in a total volume of 2 mL per well and incubated at 37°C with 5% CO_2_ for 24 hours. L-Cys@CeNPs and D-Cys@CeNPs were added at their IC_50_ concentrations, as well as at concentrations above and below the IC_50_ values, in a total volume of 2 mL of culture medium. After 24 hours of incubation, the culture medium containing non-adherent cells was carefully collected and transferred to a centrifuge tube. Adherent cells were then detached using 0.25% trypsin-EDTA solution, and the detached cells were combined with the previously collected medium to ensure comprehensive collection of both adherent and non-adherent cells. The collected cells were centrifuged at 1300 rpm for 5 minutes and resuspended in 1 mL of 1**×** DPBS containing 2% FBS. Cell counting was performed using the LUNA-II Automated Cell Counter. Subsequently, cells were centrifuged again, and the pellet was resuspended in 100 µL of Cell Staining Buffer. For apoptosis detection using Pacific Blue^TM^ Annexin V/PI Apoptosis Detection Kit (Biolegend, Cat#640928), 1 µL of Annexin V and 2 µL of PI were added to a suspension of 200,000 cells in a light-protected environment. Following a 15-minute incubation at room temperature in the dark, 100 µL of Cell Staining Buffer was added to stabilize the cells. Flow cytometry analysis was performed using a BD FACS Canto II instrument, and experiments were repeated in four or five independent sets. Data were analyzed using FlowJo V10 software. Additionally, cells treated with D- & L-Cys@CeNPs were collected for RNA isolation and stored at -80°C for subsequent PCR assays.

### 6. Gene Expression Profiling via qPCR

To investigate the potential anticancer mechanisms of D-Cys@CeNPs and L-Cys@CeNPs, gene expression profiling was performed using quantitative PCR (qPCR) to analyze the expression levels of TNFAIP3, IKBKG, and NFKBIA, with β-actin serving as the housekeeping control gene. RNA isolation and cDNA synthesis were performed prior to qPCR analysis, and the primer sequences used are listed in Table 1 of the supplementary file.

These genes were selected for their critical roles in NF-κB pathway regulation, providing insights into how CeNP treatment modulates pathway activation. TNFAIP3 (A20) acts as a negative regulator that terminates NF-κB signaling by deubiquitinating upstream components, thereby preventing prolonged activation [33]. IKBKG (NEMO) encodes an essential scaffolding protein for assembling the IKK complex, which phosphorylates IκB proteins to initiate canonical NF-κB activation [85]. NFKBIA (IκBα) functions as a feedback inhibitor, sequestering NF-κB dimers in the cytoplasm to maintain balanced signaling [86]. The selection of these genes was further guided by cell line-specific mutation profiles obtained from Cell Model Passport, COSMIC, and DepMap databases. LoVo cells carry a TNFAIP3 frameshift mutation (p.C624fs*73), potentially impairing A20 function and leading to enhanced NF-κB activation. COLO-201 cells exhibit TNFAIP3 amplification, which may strengthen its regulatory role but could also create pathway imbalances. In contrast, DLD-1 cells harbor single nucleotide variations in TNFAIP3 (p.A648S) and NFKBIA (p.P114Q), which may influence pathway activity [87–89]. These genetic distinctions provide a foundation for hypothesizing that CeNP treatments will differentially affect NF-κB signaling across these models.

#### 6.1 Total RNA Extraction and cDNA Synthesis

Total RNA was extracted using the Direct-zol™ RNA Miniprep Plus Kit (Cat#R2073, ZymoResearch). For this process, 100,000 cells were seeded per well in 6-well plates and divided into two groups: control and treated. After 24 hours of incubation at 37°C with 5% CO_2_, the treated group was exposed to nanoparticles at their respective IC_50_ concentrations and incubated for another 24 hours under the same conditions. Subsequently, both adherent and non-adherent cells were collected. Non-adherent cells were first removed by collecting the culture medium, and adherent cells were detached using trypsinization. After combining both cell populations, the cells were centrifuged, washed twice with cold PBS, and TRI-Reagent was added to the pellet. RNA isolation was carried out according to the manufacturer’s protocol. The quality and concentration of RNA were determined by measuring the 260/280 nm ratio using NanoDrop™ One/OneC (Thermo Scientific, Waltham, MA, USA). RNA integrity was assessed through agarose gel electrophoresis, and samples were stored at -80°C until further use. For cDNA synthesis, 1 µg of total RNA was used with the iScript cDNA Synthesis Kit (BioRad Laboratories, CA, USA, Cat# 1708891). The reaction components and volumes for cDNA synthesis are detailed in Supplementary Table 2. The total reaction volume, including oligo(dT) and random primers, was set to 20 µL. cDNA synthesis was carried out in a thermal cycler according to the protocol outlined in Supplementary Table 3.

#### 6.2. RT-qPCR Analysis

cDNA samples, diluted to a uniform concentration of 10 ng/µL, were amplified using the SYBR Green PCR Master Mix (Bio-Rad, Cat# 1725120). Each 20 µL reaction contained 40 ng of cDNA, 5 µM forward and reverse primers, reaction mix, and nuclease-free water. Real-time PCR was carried out on the AriaMx Real-Time PCR System (Agilent, USA) under the following conditions: an initial denaturation at 95°C for 2 minutes, followed by 40 cycles of 95°C for 10 seconds, 58°C for 30 seconds, and 72°C for 30 seconds. Each sample was analyzed with three biological replicates. Gene expression levels were quantified relative to β-actin, which served as the housekeeping gene. The 2⁻^ΔΔCt^ method was used to determine the relative mRNA expression of target genes. The amplification curves and reaction component protocols are presented in Supplementary Tables 4 and 5.

## STATISTICAL ANALYSIS

All data were expressed as mean ± standard deviation (mean ± SD). Statistical analyses were performed using GraphPad PRISM software. Differences between groups were assessed using one-way and two-way ANOVA, followed by Tukey’s post hoc test to determine significance between nanoparticle-treated and control groups. Statistical significance levels were reported as follows: *p < 0.05, **p < 0.01, ***p < 0.001, and ****p < 0.0001. Non-significant differences were indicated as “ns.”

## SUPPORTING INFORMATION

Cytotoxicity results of each cell line

TNFR signaling pathway

Figure SI1-14

Table SI1-13

## ACKNOWLEDGEMENT

E.S.T.E. and A.D. wrote the manuscript. E.S.T.E. synthesized and characterized the nanoparticles, analyzed the data, prepared the figures. A.D. performed the cell culture experiments and prepared the corresponding figures. E.S.T.E. made further analysis on cell culture experiments as needed. S.A. performed the qPCR experiments and analyzed the qPCR data. B.K. and N.S. assisted with nanoparticle synthesis and characterization and contributed to data interpretation. N.S. contributed to manuscript editing. A.M.Y and A.D. performed the flow cytometry experiments and supported flow cytometry data analysis. H.Y. and N.A.K. conceived and supervised the project.

The authors acknowledge the University of Michigan College of Engineering for financial support and the Michigan Center for Materials Characterization for the use of TEM and XPS. This study is supported by the International Centre for Genetic Engineering and Biotechnology (ICGEB) under CRP/TUR 18-03 project number, TUBA-GEBIP Distinguished Young Scientist Award (Hilal Yazici, 2019), Turkish Scientific and Technical Research Council (TÜBITAK) under the KAMAG-1007 Project number: 113G100.

## Declaration of Generative AI and AI-assisted technologies in the writing process

During the preparation of this work, the authors used ChatGPT to improve the language, flow, and clarity of the manuscript and also utilized Gemini 3.1 to enhance the artistic quality of some figures. All AI tool usage is clearly stated in the figure legends, if applicable. After using these tools, the authors reviewed and edited the content as needed and take full responsibility for its accuracy.

