## Supplementary material for "Enantiomer-Dependent Biological Activity of Cysteine-Coated Ceria Nanoparticles in Colorectal Cancer Cells": SI

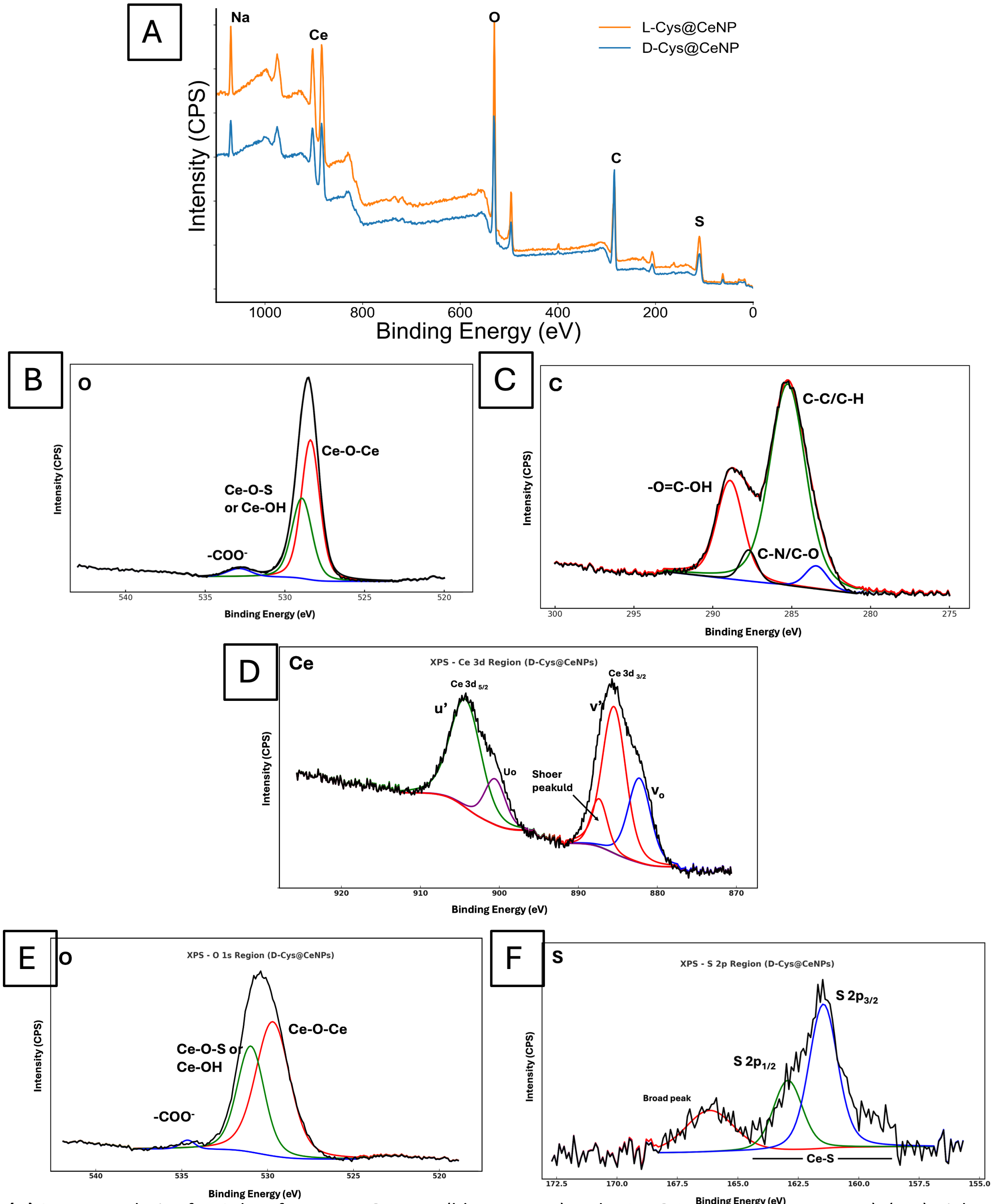

**Figure SI 1: (A)** Survey analysis of XPS data for D –Cys@CeNPs (blue spectra) and L-Cys@CeNPs (orange spectra). (B-F) High-resolution XPS spectra of L-Cys@CeNP and D-Cys@CeNP nanoparticles**. (B) O** 1s spectrum of L-Cys@CeNP shows deconvoluted peaks corresponding to Ce–O–Ce (lattice oxygen), Ce–OH or Ce–O–S (surface groups), and carboxylates ($–COO^–$), indicating mixed oxide and ligand contributions. (**C)** C 1s spectrum of L-Cys@CeNP reveals components attributed to C–C/C–H (hydrocarbon backbone), C–OH, C–N/C–O, and carboxylic functionalities, confirming cysteine surface coating. **(D)** Ce 3d region of D-Cys@CeNP displays characteristic $Ce^{3+}$ features, including u′, $u_0$, v′, and $v_0$ peaks. The absence of $Ce^{4+}$ satellite peaks confirms a predominantly $Ce^{3+}$ oxidation state. (**E)** O 1s spectrum of D-Cys@CeNP shows similar features to (A), with peaks for Ce–O–Ce, Ce–OH, and $–COO^–$, supporting the presence of cerium oxide and surface ligands. **(F)** S 2p spectrum of D-Cys@CeNP exhibits distinct peaks for S $2p_{3/2}$ and S $2p_{1/2}$ (~163–165 eV), assigned to thiol-bound sulfur (Ce–S), along with a broad peak suggesting oxidized sulfur species or disulfide/thiol variants.

### Time- and Dose-Dependent Cytotoxicity and Therapeutic Selectivity of D-Cys@CeNPs and L-Cys@CeNPs on Cell Lines

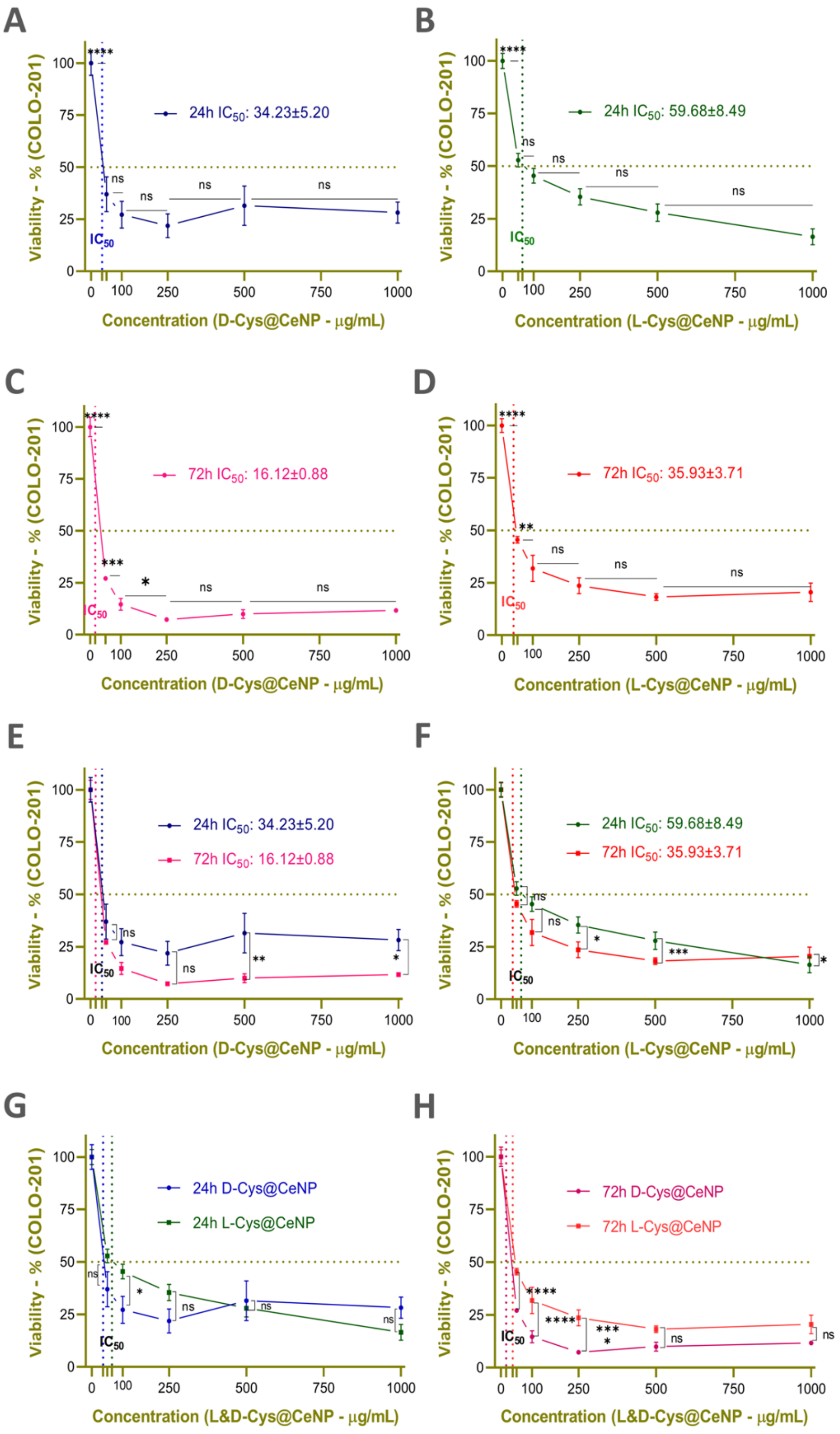

**Figure SI 2. Cytotoxic effects of D-Cys@CeNP and L-Cys@CeNP on COLO-201 cells. (A & B):** Cell viability of COLO-201 cells treated with D-Cys@CeNP **(A)** and L-Cys@CeNP **(B)** for 24 h at increasing concentrations. **(C & D):** Cell viability of COLO-201 cells treated with D-Cys@CeNP **(C)** and L-Cys@CeNP **(D)** for 72 h at increasing concentrations. **(E & F)** Comparison of 24 h and 72 h treatments for D-Cys@CeNP **(E)** and L-Cys@CeNP **(F). (G & H)**Comparison of D-Cys@CeNP and L-Cys@CeNP treatments at 24 h **(G)** and 72 h **(H).** The results represent the mean ± SD of three independent biological replicates (n=3). The dotted vertical lines indicate the $IC_{50}$ values, which are also displayed in the figure. Statistical significance was analyzed using one-way ANOVA followed by Tukey's post-hoc test: (ns = non-significant, * $p < 0.05$, ** $p < 0.01$, *** $p < 0.001$, **** $p < 0.0001$).

**COLO-201 Cells**

Cytotoxicity analyses revealed that the effects of D-Cys@CeNPs and L-Cys@CeNPs on COLO-201 cell viability were both time- and dose-dependent. **(Figure 3-pink and SI 2)**. COLO-201 cells were highly sensitive to nanoparticle treatment.

24-hour exposure: D-Cys@CeNPs induced a sharp reduction in cell viability, particularly at lower doses (50–250 μg/mL), while at higher concentrations (500–1000 μg/mL), the cytotoxic effect appeared to plateau **(Figure 3A-pink and SI 2A**). In contrast, L-Cys@CeNPs exhibited a more gradual dose-dependent decrease in viability, suggesting a controlled apoptotic response **(Figure 3B- pink and SI 2B).**

72-hour exposure: Prolonged treatment significantly enhanced cytotoxicity for both enantiomers across all concentrations **(Figure 3C,D-pink and SI 2C,D)**. Notably, COLO-201 cells became more susceptible to D-Cys@CeNP-induced cytotoxicity over time, with an accelerated decline in viability at lower doses **(Figure 3C-pink and SI 2C).** Similarly, L-Cys@CeNP-treated cells exhibited a progressive but more moderate increase in cytotoxicity (Figure 3D-pink and SI 2D).

Time-dependent comparisons further reinforced these observations (Figure 3 E and Figure SI 2E&F). After 24 hours, D-Cys@CeNPs induced a rapid and pronounced cytotoxic effect, whereas L-Cys@CeNPs displayed a gradual and controlled reduction in viability **(Figure SI 2G).** After 72 hours, the cytotoxic effects of both nanoparticles significantly increased (**Figure 3 E and Figure SI 2E&F).** However, D-Cys@CeNPs remained more effective at lower concentrations, indicating sustained cytotoxic activity, while L-Cys@CeNPs maintained a more progressive yet consistent reduction in cell viability over time **(Figure SI 2H).** When comparing the enantiomer-specific responses **(Figure 3E and SI 2G-H),** at 24 hours, D-Cys@CeNPs induced a faster, more aggressive cytotoxic response, likely attributed to their to it entery the cells faster. By 72 hours, however, both nanoparticles exhibited comparable levels of cytotoxicity at higher concentrations, suggesting that prolonged exposure may equilibrate the differences between the two enantiomers. $IC_{50}$ calculations showed that COLO-201 cells were highly sensitive to both nanoparticle formulations, with a notably stronger response to D-Cys@CeNPs (Figure 3E and Figure SI 2E-H)**.** At 24 hours, the $IC_{50}$ values were $34.23 \pm 5.20$ μg/mL for D-Cys@CeNPs and $59.68 \pm 8.49$ μg/mL for L-Cys@CeNPs. By 72 hours, these values decreased significantly to $16.12 \pm 0.88$ μg/mL and $35.93 \pm 3.71$ μg/mL, respectively, indicating time-dependent cytotoxic enhancement ($p < 0.01$). These results highlight the importance of both treatment duration and nanoparticle chirality in modulating therapeutic outcomes in CRC.

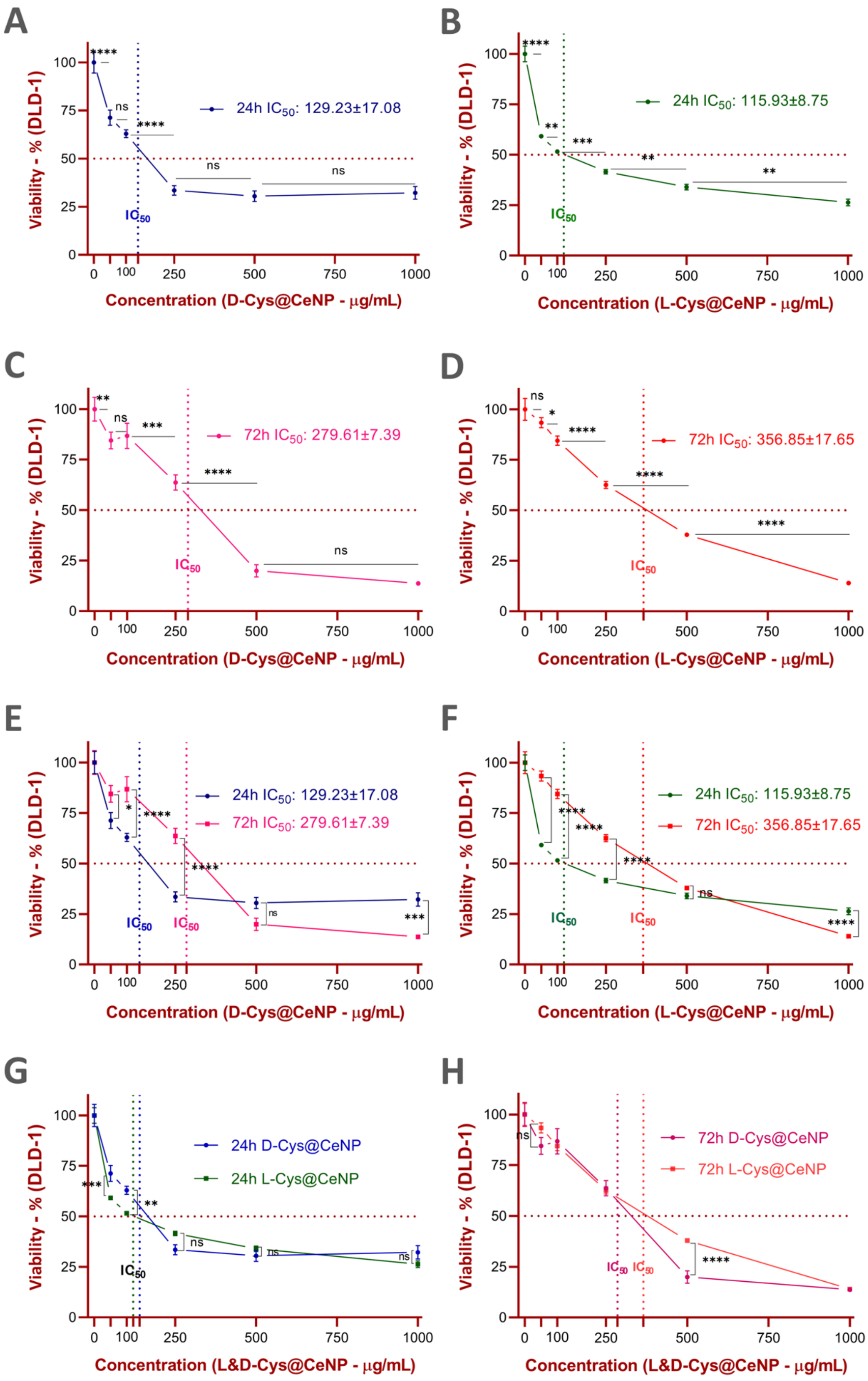

**Figure SI 3.** Cytotoxic effects of D-Cys@CeNP and L-Cys@CeNP on DLD-1 cells. DLD-1 colorectal cancer cells were treated with D-Cys@CeNP and L-Cys@CeNP at concentrations of 0 (control), 50, 100, 250, 500, and 1000 µg/mL for 24 h and 72 h. Cell viability was assessed using the WST-1 reagent, and the control group (0 µg/mL) was set as 100% viability. The results represent the mean ± SD of three independent biological replicates (n=3). Panel descriptions: (A & B): Cell viability of DLD-1 cells treated with D-Cys@CeNP (A) and L-Cys@CeNP (B) for 24 h at increasing concentrations. (C & D): Cell viability of DLD-1 cells treated with D-Cys@CeNP (C) and L-Cys@CeNP (D) for 72 h at increasing concentrations. (E & F): Comparison of 24 h and 72 h treatments for D-Cys@CeNP (E) and L-Cys@CeNP (F). (G & H): Comparison of D-Cys@CeNP and L-Cys@CeNP treatments at 24 h (G) and 72 h (H).

**DLD-1 Cells**

Cytotoxicity analyses in DLD-1 cells also demonstrated that both D-Cys@CeNPs and L-Cys@CeNPs exhibited dose- and time-dependent effects on cell viability **(Figure 3-green and SI 3).**

24-hour exposure**:** D-Cys@CeNPs induced a sharp reduction in viability, with a 71% survival rate at 50 µg/mL**,** and to 33% at 250 µg/mL (Figure 3A-green, Figure SI 3A). In contrast, L-Cys@CeNPs caused an earlier, more gradual reduction in viability with 59% survival at 50 µg/mL (Figure 3B-green, Figure SI 3B), indicating a slower yet steady onset of cell death.

72-hour exposure: Both nanoparticles exhibited stronger cytotoxicity, with D-Cys@CeNPs further reducing viability to 19% at 500 µg/mL and 13% at 1000 µg/mL**. (**Figure 3 C-green, SI 3C**)** L-Cys@CeNPs showed a more moderate decline in viability at lower doses but still demonstrated a dose-dependent reduction in viability (Figure 3D-green, SI 3D).

In the time-dependent comparison (Figure SI 3E&F), both nanoparticles showed a clear increase in cytotoxicity over time. D-Cys@CeNP-treated cells displayed a steeper drop in viability over time, whereas L-Cys@CeNPs exhibited a more consistent and controlled decline. When D-Cys@CeNPs and L-Cys@CeNPs were directly compared at each time point (Figure 3E and SI 3G-H), at 24 hours, D-Cys@CeNPs caused a more immediate cytotoxic effect, whereas L-Cys@CeNPs led to a more progressive decline. By 72 hours, both nanoparticles exhibited strong cytotoxicity, particularly at higher concentrations, suggesting a cumulative effect over prolonged exposure.

$IC_{50}$ calculations confirmed that DLD-1 cells were moderately responsive to both nanoparticle formulations, with comparable sensitivity to D-Cys@CeNPs and L-Cys@CeNPs at early time points (Figure 3E; SI 3E–H). At 24 hours, the $IC_{50}$ values were 129.23 ± 17.08 µg/mL for D-Cys@CeNPs and 115.93 ± 8.75 µg/mL for L-Cys@CeNPs. By 72 hours, these values increased significantly to 279.61 ± 7.39 µg/mL and 356.85 ± 17.65 µg/mL, respectively, suggesting a reduced cytotoxic response with prolonged exposure.

These findings point to a potential adaptive resistance mechanism in DLD-1 cells—possibly involving NF-κB-mediated survival signaling or enhanced antioxidant capacity—that limits the long-term efficacy of cerium oxide nanoparticles. The differences in early versus late response further emphasize the role of nanoparticle chirality and treatment duration in shaping therapeutic outcomes in inflammation-associated CRC subtypes.

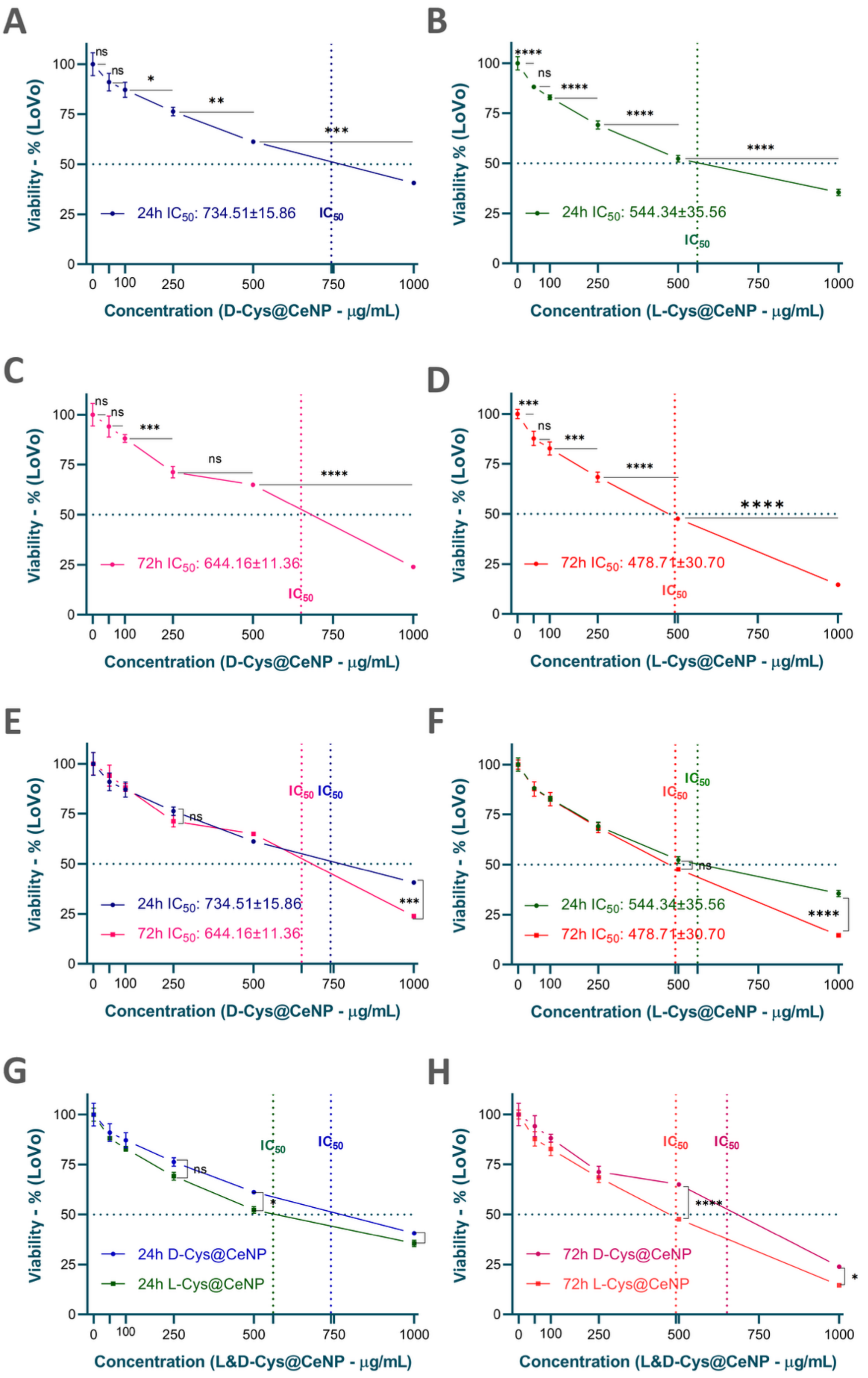

**Figure SI 4.** Cytotoxic effects of D-Cys@CeNP and L-Cys@CeNP on LoVo cells.. Panel descriptions: **(A & B)**: Cell viability of LoVo cells treated with D-Cys@CeNP **(A)** and L-Cys@CeNP **(B)** for 24 h at increasing concentrations. (**C & D**): Cell viability of LoVo cells treated with D-Cys@CeNP **(C)** and L-Cys@CeNP **(D)** for 72 h at increasing concentrations. **(E & F):** Comparison of 24 h and 72 h treatments for D-Cys@CeNP **(E)** and L-Cys@CeNP **(F).** (**G & H):** Comparison of D-Cys@CeNP and L-Cys@CeNP treatments at 24 h **(G)** and 72 h **(H).** The dotted vertical lines indicate the $IC_{50}$ values, which are also displayed in the figure.Statistical significance was analyzed using one-way ANOVA followed by Tukey's post-hoc test:(ns = non-significant, * $p < 0.05$, ** $p < 0.01$, *** $p < 0.001$, **** $p < 0.0001$).

**LoVo Cells**

Cytotoxicity analyses on LoVo cells revealed that this cell line exhibited greater resistance to D-Cys@CeNPs, whiles L-Cys@CeNPs demonstrated a stronger cytotoxic effect over time (Figure 3-purple, SI 4). However, overall, LoVo cells were more resistant to both nanoparticles compared to other cell lines.

24-hour exposure**:** D-Cys@CeNPs showed a moderate cytotoxic effect at 500–1000 µg/mL**,** with viability rates at 61% at 500 µg/mL and 40% at 1000 µg/mL (Figure 3A-purple, SI 4A). In contrast, L-Cys@CeNPs exhibited a more pronounced early-stage cytotoxic effect**,** reducing viability to 52% at 500 µg/mL and 35% at 1000 µg/mL (Figure 3B-purple, SI 4B).

72-hour exposure**:** LoVo cells continued to show considerable resistance to D-Cys@CeNPs at lower doses, but viability decreased to 23% at 1000 µg/mL, confirming an increased cytotoxic effect over time (Figure 3C-purple, SI 4C). L-Cys@CeNPs showed a pronounced effect at 72 hours, reducing viability to 14% at 1000 µg/mL (Figure 3D-purple, SI 4D).

In the time-dependent comparison (Figure SI 4E&F), both nanoparticles exhibited time-dependent cytotoxicity, but L-Cys@CeNPs produced a more pronounced long-term effect. When directly comparing the enantiospecific effects of D-Cys@CeNPs and L-Cys@CeNPs (Figure 3E and SI 4G-H), at 24 hours, L-Cys@CeNPs exhibited a steeper decline in viability, while D-Cys@CeNPs maintained a more gradual effect. By 72 hours, both nanoparticles demonstrated significant cytotoxicity at higher concentrations, with L-Cys@CeNPs maintaining a stronger effect overall.

$IC_{50}$ calculations confirmed that LoVo cells were relatively resistant to both D-Cys@CeNPs and L-Cys@CeNPs, with higher $IC_{50}$ values observed at both time points compared to COLO-201 and DLD-1 cells (Figure 3E; SI 4E–H). At 24 hours, the $IC_{50}$ was 734.51 ± 15.86 µg/mL for D-Cys@CeNPs and 544.34 ± 35.56 µg/mL for L-Cys@CeNPs. After 72 hours, the values decreased modestly to 644.16 ± 11.36 µg/mL and 478.71 ± 30.70 µg/mL, respectively, indicating a limited but measurable increase in cytotoxic effect with extended exposure.

These results suggest that LoVo cells possess robust antioxidant defenses or stress-adaptive mechanisms that confer resistance to CeNP-induced cytotoxicity. The consistently stronger effect of L-Cys@CeNPs compared to D-Cys@CeNPs—particularly over time—emphasizes the relevance of chirality in modulating therapeutic response in aggressive, redox-resistant CRC models. Particles may not enter the cells.

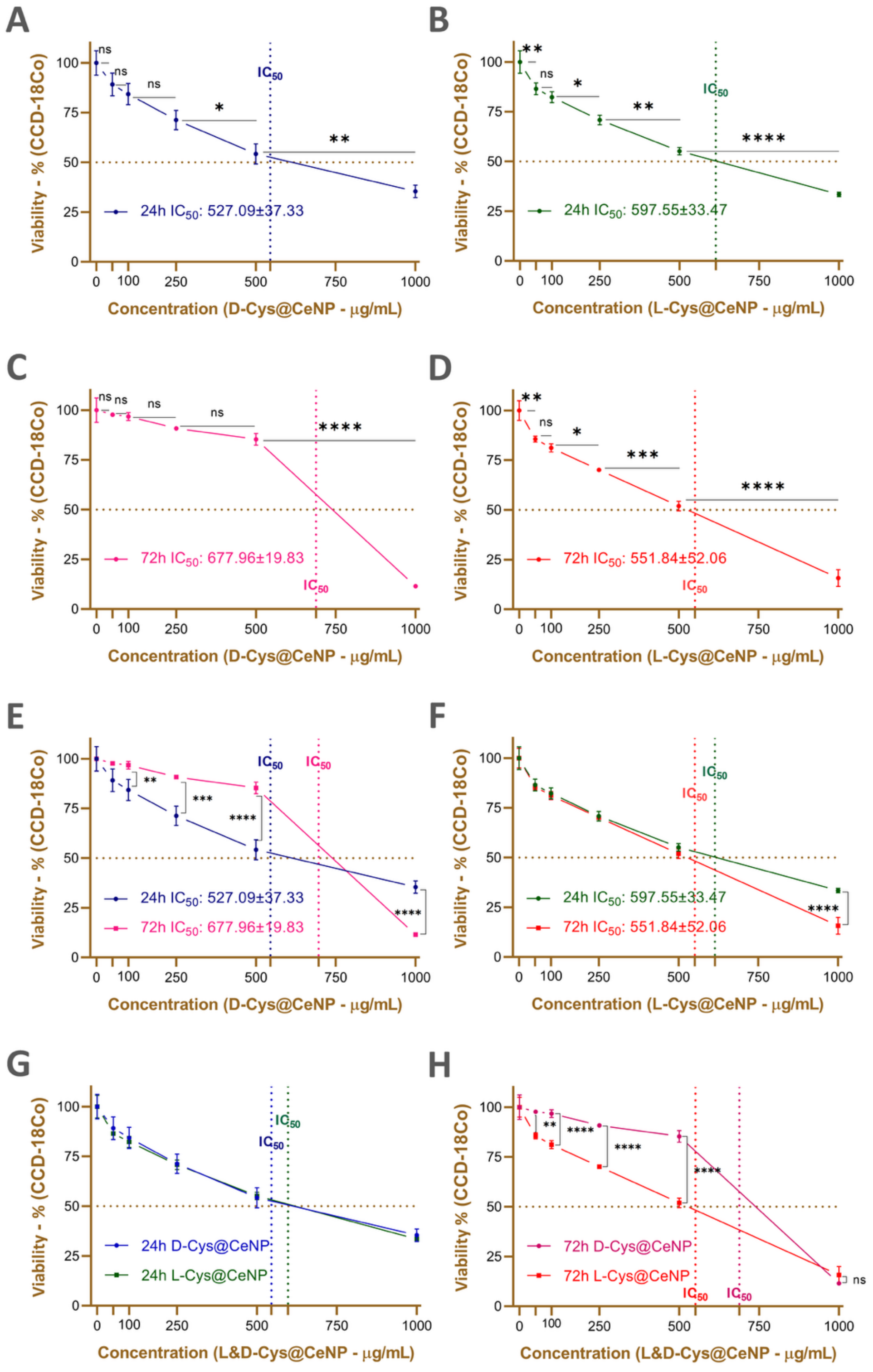

**Figure SI 5. Cytotoxic effects of D-Cys@CeNP and L-Cys@CeNP on CCD-18Co cells (A & B)** Cell viability of CCD-18Co cells treated with D-Cys@CeNP **(A)** and L-Cys@CeNP **(B)** for 24 h at increasing concentrations. **(C & D)** Cell viability of CCD-18Co cells treated with D-Cys@CeNP **(C)** and L-Cys@CeNP (D) for 72 h at increasing concentrations**. (E & F):** Comparison of 24 h and 72 h treatments for D-Cys@CeNP **(E)** and L-Cys@CeNP **(F). (G & H**): Comparison of D-Cys@CeNP and L-Cys@CeNP treatments at 24 h **(G)** and 72 h **(H).** The results represent the mean ± SD of three independent biological replicates (n=3). The dotted vertical lines indicate the $IC_{50}$ values, which are also displayed in the figure.Statistical significance was analyzed using one-way ANOVA followed by Tukey's post-hoc test: (ns = non-significant, * $p < 0.05$, ** $p < 0.01$, *** $p < 0.001$, **** $p < 0.0001$).

**CCD-18Co Cells**

Cytotoxicity analyses revealed that CCD-18Co healthy colon fibroblasts exhibited higher resistance to both D-Cys@CeNPs and L-Cys@CeNPs compared to cancer cells, with cell viability being significantly maintained at lower doses but gradually decreasing at higher concentrations over time (Figure 3-blue, SI 5A-H).

24-hour exposure: D-Cys@CeNP-treated CCD-18Co cells maintained over 90% viability at lower doses (50–250 µg/mL), but dropped to approximately 52% at 500 µg/mL and 32% at 1000 µg/mL (Figure 3A-blue, SI 5A). Similarly, L-Cys@CeNP-treated cells retained high viability at lower doses, but at 500 µg/mL, viability dropped to approximately 54%, and at 1000 µg/mL, it further declined to around 33% (Figure 3B-blue SI 5B).

72-hour exposure**:** An interesting trend was observed for D-Cys@CeNPs, where cell viability at 500 µg/mL was better preserved compared to 24 hours, reaching approximately 82%. This suggests an adaptive response or recovery at moderate concentrations (Figure 3C-blue, SI 5C). In contrast, L-Cys@CeNP-treated cells exhibited a similar viability at 500 µg/mL over both time points, maintaining around 50%. At 1000 µg/mL, both nanoparticles showed a more substantial cytotoxic effect, with viability decreasing to 11% for D-Cys@CeNPs and 19% for L-Cys@CeNPs (Figure 3D-blue, SI 5D).

The time-dependent analysis (Figure SI 5E&F) revealed that both nanoparticles exhibited gradual cytotoxicity, though CCD-18Co cells remained significantly more resistant compared to cancer cell lines. D-Cys@CeNP-treated cells adapted better over time at all concentrations except for 1000 µg/mL, where the cytotoxicity became more pronounced. L-Cys@CeNPs induced a more pronounced decrease over time, but no drastic viability loss was observed.

When comparing the enantiospecific effects **(Figure 3E, SI 5G&H)**, CCD-18Co cells maintained higher viability against both nanoparticles compared to cancer cells. Notably, D-Cys@CeNP-treated cells exhibited a stronger adaptive response at 500 µg/mL over time, maintaining higher viability compared to 24 hours. In contrast, L-Cys@CeNP-treated cells showed a relatively stable response at 500 µg/mL across both time points. However, at higher concentrations and prolonged exposure, a dose-dependent cytotoxic effect became evident.

$IC_{50}$ calculations confirmed that CCD-18Co normal colon fibroblasts were more resistant to both D-Cys@CeNPs and L-Cys@CeNPs compared to all CRC cell lines (Figure 3E, SI 5E–H). At 24 hours, the $IC_{50}$ was 527.09 ± 37.33 µg/mL for both D-Cys@CeNPs and L-Cys@CeNPs. After 72 hours, the $IC_{50}$ values increased to 677.96 ± 19.83 µg/mL for D-Cys@CeNPs and 551.84 ± 52.06 µg/mL for L-Cys@CeNPs, indicating minimal long-term cytotoxicity in normal cells and a relatively high threshold for toxicity.

These findings suggest that CCD-18Co cells possess effective protective mechanisms against oxidative stress and remain largely unaffected by CeNP exposure at lower to moderate concentrations. The increased $IC_{50}$ for D-Cys@CeNPs at 72 hours may reflect a delayed adaptive response that helps preserve cell viability. The overall higher $IC_{50}$ values and sustained viability underscore the favorable safe profile of both nanoparticle formulations, particularly in normal tissue contexts.

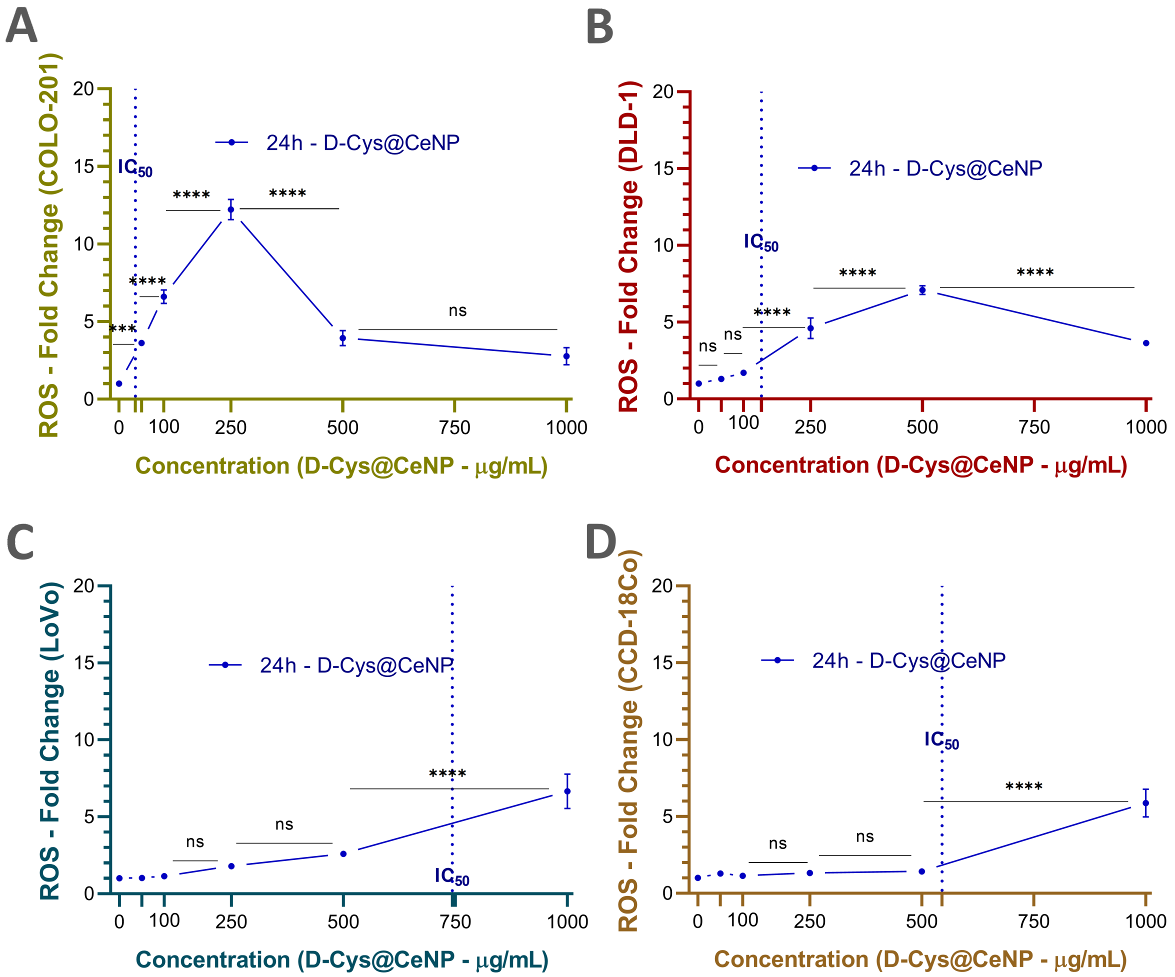

**Figure SI 6. ROS generation in colorectal cancer and normal fibroblast cells treated with D-Cys@CeNP for 24 hours**. **(A & B)** COLO-201 cells treated with D-Cys@CeNP for 24 h. **(A)** Cell viability (%); **(B)** ROS fold change. **(C & D)** DLD-1 cells treated with D-Cys@CeNP for 24 h. **(C)** Cell viability; **(D)** ROS fold change**. (E & F)**: LoVo cells treated with D-Cys@CeNP for 24 h. **(E)** Cell viability; (F) ROS fold change. **(G & H)**: CCD-18Co cells treated with D-Cys@CeNP for 24 h. **(G)** Cell viability; **(H)** ROS fold change. The results represent the mean ± SD of three independent biological replicates (n=3). The vertical dashed lines indicate the $IC_{50}$ values on the X-axis. Statistical significance was analyzed using one-way ANOVA followed by Tukey's post-hoc test: (ns = non-significant, * $p < 0.05$, ** $p < 0.01$, *** $p < 0.001$, **** $p < 0.0001$).

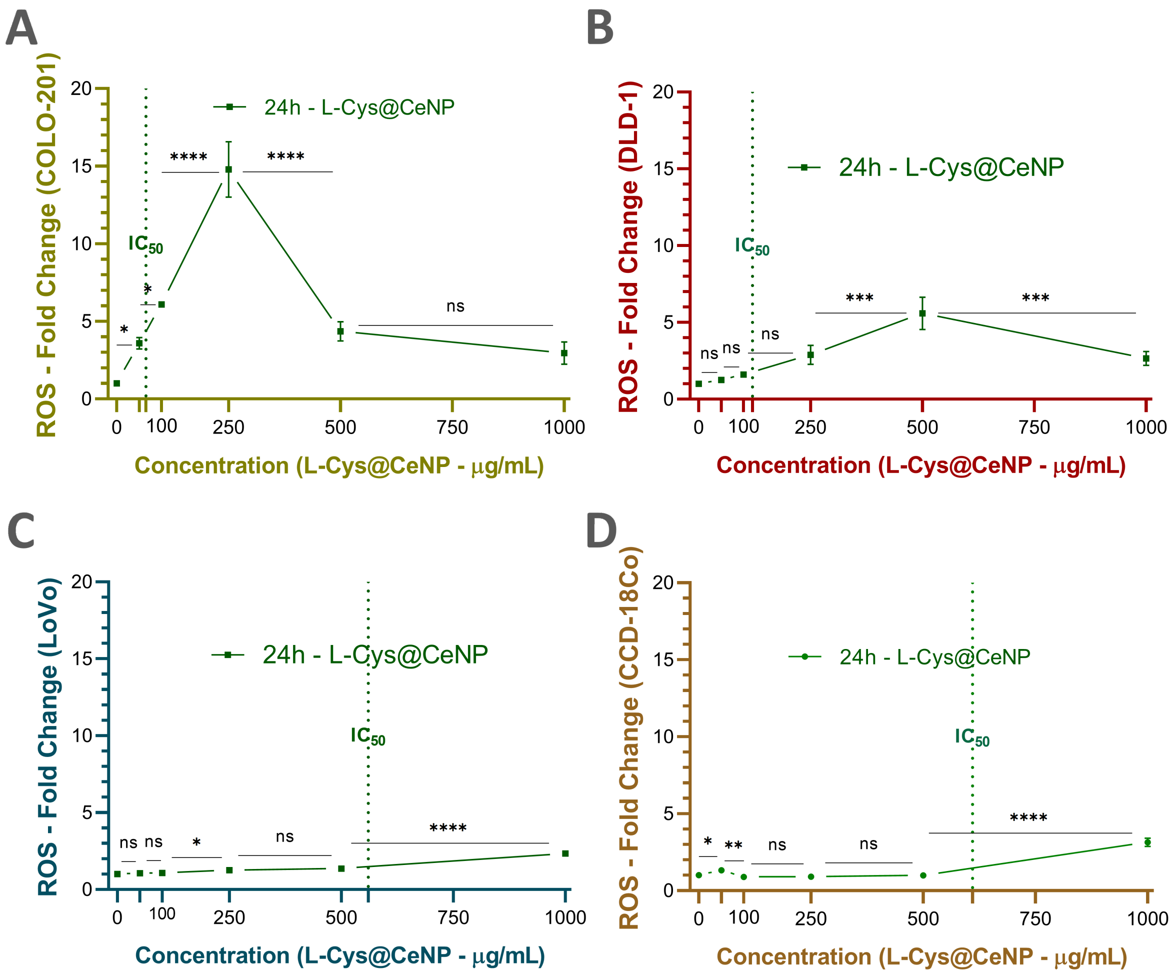

**Figure SI7. ROS generation in colorectal cancer and normal fibroblast cells treated with L-Cys@CeNP for 24 hours. (A & B)** COLO-201 cells treated with L-Cys@CeNP for 24 h. **(A)** Cell viability (%); **(B)** ROS fold change. **(C & D)**: DLD-1 cells treated with L-Cys@CeNP for 24 h. **(C)** Cell viability; **(D)** ROS fold change. **(E & F):** LoVo cells treated with L-Cys@CeNP for 24 h. **(E)** Cell viability; **(F)** ROS fold change. **(G & H)** CCD-18Co cells treated with L-Cys@CeNP for 24 h. **(G)** Cell viability; **(H)** ROS fold change. The results represent the mean ± SD of three independent biological replicates (n=3). The vertical dashed lines indicate the $IC_{50}$ values on the X-axis. Statistical significance was analyzed using one-way ANOVA followed by Tukey's post-hoc test: (ns = non-significant, * $p < 0.05$, ** $p < 0.01$, *** $p < 0.001$, **** $p < 0.0001$).

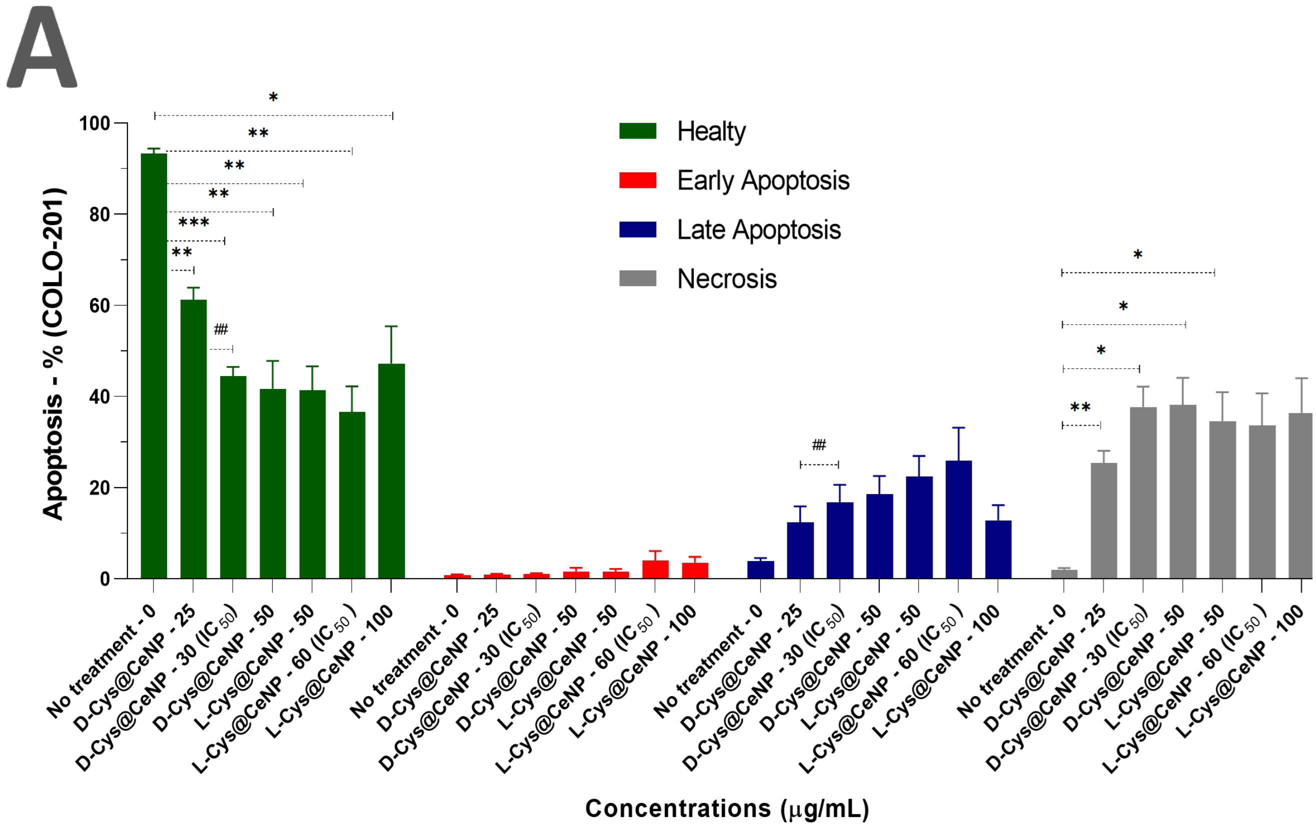

**Figure SI8. Flow cytometry-based apoptosis analysis of COLO-201 cells treated with D-Cys@CeNP and L-Cys@CeNP for 24 hours.** Panel descriptions: **(A):** Quantitative bar graphs showing the mean ± SD percentages of healthy, early apoptotic, late apoptotic, and necrotic cells across five biological replicates (n=5). Statistical comparisons between treatment groups are indicated. **(**The results represent the mean ± SD of five independent biological replicates (n=5). Statistical significance was analyzed using two-way ANOVA followed by Tukey's post-hoc test: (ns = non-significant, * $p < 0.05$, ** $p < 0.01$, *** $p < 0.001$, **** $p < 0.0001$).

**Table SI1:** The percentage distribution of healthy cells, early apoptosis, late apoptosis, and necrosis in COLO-201 colon cancer cells treated with L&D@CeNPs (n=5).

| **COLO-201 Colon Cancer Cells** (NPs Conc. - µg/mL) | **Healty** (Mean± SD - %) | **Early Apoptosis** (Mean± SD - %) | **Late Apoptosis** (Mean± SD - %) | **Necrosis** (Mean± SD - %) |
|---|---|---|---|---|
| **No treatment - 0** | 93.32 ± 2.43 | 0.80 ± 0.36 | 3.92 ± 1.49 | 1.97 ± 0.92 |
| **D-Cys@CeNP - 25** | 61.24 ± 6.00 | 0.97 ± 0.26 | 12.37 ± 7.92 | 25.44 ± 5.98 |
| **D-Cys@CeNP - 30 ($IC_{50}$)** | 44.44 ± 4.58 | 1.04 ± 0.44 | 16.82 ± 8.54 | 37.68 ± 10.04 |
| **D-Cys@CeNP - 50** | 41.68 ± 13.83 | 1.61 ± 1.89 | 18.55 ± 8.97 | 38.16 ± 13.28 |
| **L-Cys@CeNP - 50** | 41.36 ± 11.75 | 1.57 ± 1.26 | 22.46 ± 10.09 | 34.62 ± 14.13 |
| **L-Cys@CeNP - 60 ($IC_{50}$)** | 36.60 ± 12.52 | 3.94 ± 4.76 | 25.88 ± 16.34 | 33.60 ± 15.94 |
| **L-Cys@CeNP - 100** | 47.22 ± 18.30 | 3.56 ± 2.79 | 12.81 ± 7.53 | 36.40 ± 17.05 |

A

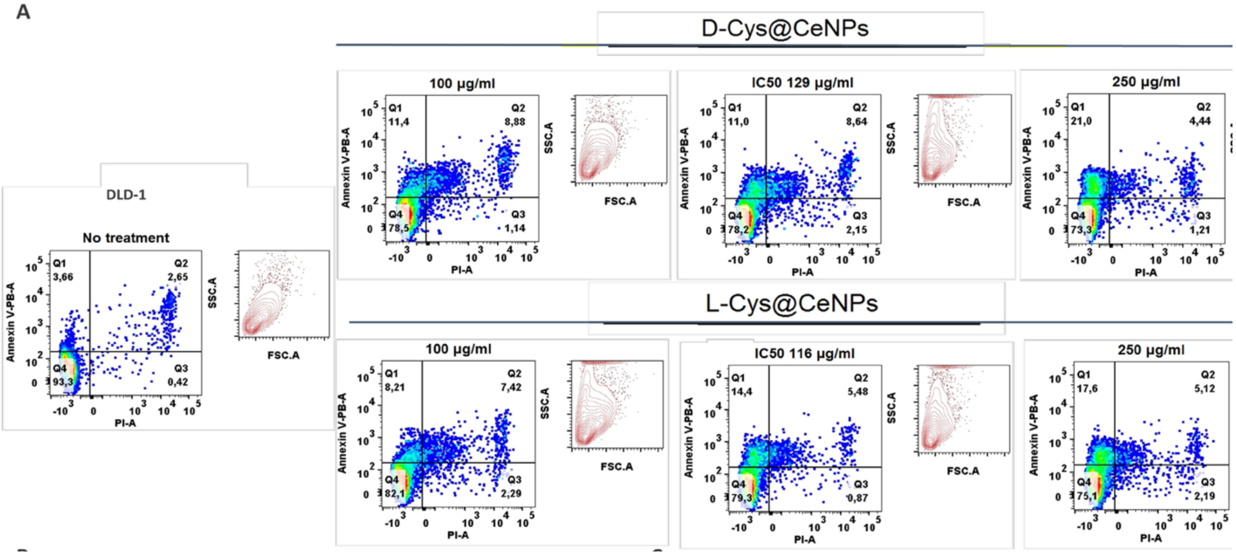

B

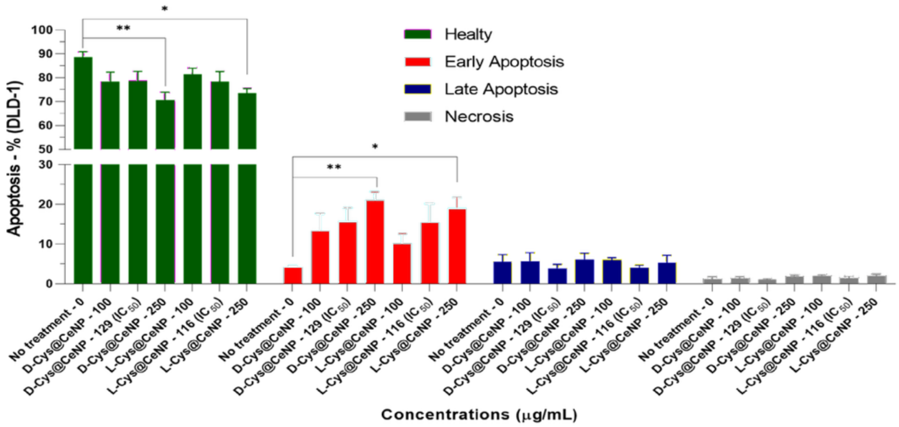

**Figure SI9**. **Flow cytometry-based apoptosis analysis of DLD-1 cells treated with D-Cys@CeNP and L-Cys@CeNP for 24 hours. (A)** Representative flow cytometry plots for each treatment condition in DLD-1 cells. Quadrant analysis distinguishes healthy (Q4; Annexin V-/PI-), early apoptotic (Q1; Annexin V+/PI-), late apoptotic (Q2; Annexin V+/PI+), and necrotic (Q3; Annexin V-/PI+) populations. **(B)** Quantitative bar graphs showing the mean ± SD percentages of healthy, early apoptotic, late apoptotic, and necrotic cells across five biological replicates (n=5). Statistical comparisons between treatment groups are indicated. The results represent the mean ± SD of five independent biological replicates (n=5). Statistical significance was analyzed using two-way ANOVA followed by Tukey's post-hoc test: (ns = non-significant, * $p < 0.05$, ** $p < 0.01$, *** $p < 0.001$, **** $p < 0.0001$).

**Table SI2:** The percentage distribution of healthy cells, early apoptosis, late apoptosis, and necrosis in DLD-1 colon cancer cells treated with L&D@CeNPs (n=5).

| **DLD-1 Colon Cancer Cells** (NPs Conc. - µg/mL) | **Healty** (Mean± SD - %) | **Early Apoptosis** (Mean± SD - %) | **Late Apoptosis** (Mean± SD - %) | **Necrosis** (Mean± SD - %) |
|---|---|---|---|---|
| No treatment - 0 | 88.68 ± 4.96 | 4.17 ± 0.72 | 5.55 ± 4.05 | 1.22 ± 1.14 |
| D-Cys@CeNP - 100 | 78.26 ± 9.19 | 13.24 ± 9.87 | 5.66 ± 4.87 | 1.43 ± 0.68 |
| D-Cys@CeNP - 129 ($IC_{50}$) | 78.62 ± 9.02 | 15.66 ± 7.87 | 3.91 ± 2.21 | 1.09 ± 0.47 |
| D-Cys@CeNP - 250 | 70.60 ± 7.50 | 21.04 ± 4.68 | 6.09 ± 3.68 | 1.83 ± 0.79 |
| L-Cys@CeNP - 100 | 81.42 ± 5.55 | 10.15 ± 5.43 | 5.99 ± 1.32 | 2.10 ± 0.42 |
| L-Cys@CeNP - 116 ($IC_{50}$) | 78.32 ± 9.56 | 15.31 ± 10.78 | 4.07 ± 1.30 | 1.50 ± 0.90 |
| L-Cys@CeNP - 250 | 73.48 ± 4.36 | 18.96 ± 6.16 | 5.29 ± 4.23 | 2.04 ± 0.58 |

A

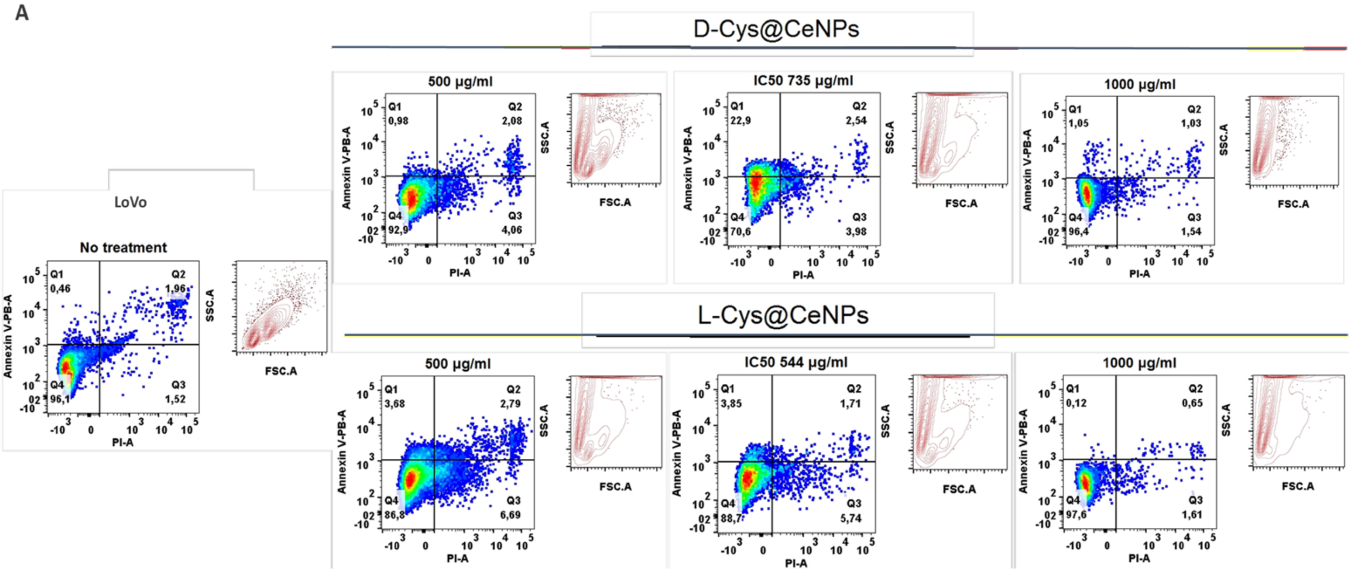

B

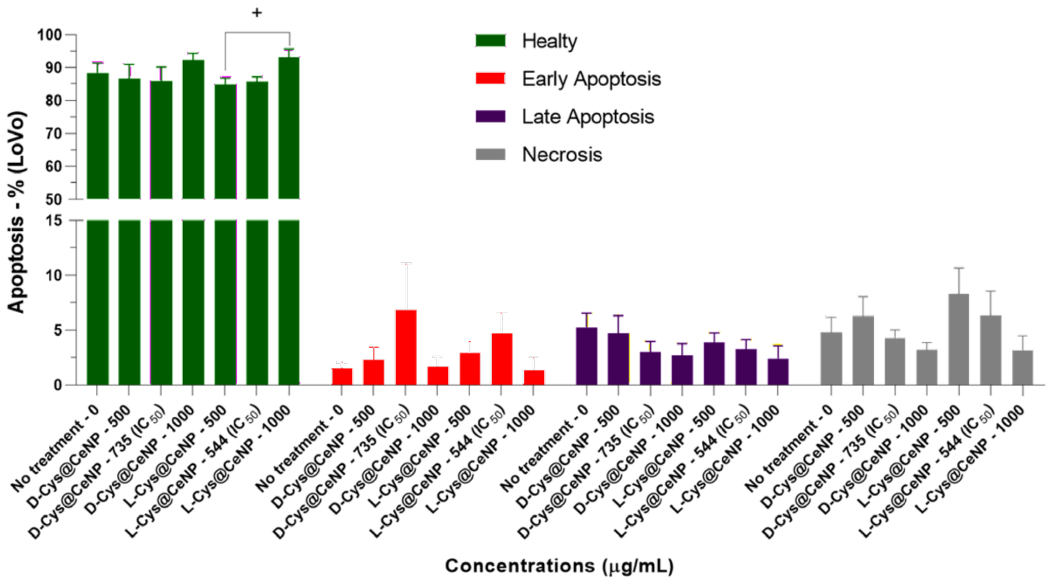

**Figure SI10**. **Flow cytometry-based apoptosis analysis of LoVo cells treated with D-Cys@CeNP and L-Cys@CeNP for 24 hours. (A)** Representative flow cytometry plots for each treatment condition in LoVo cells. Quadrant analysis distinguishes healthy (Q4; Annexin V-/PI-), early apoptotic (Q1; Annexin V+/PI-), late apoptotic (Q2; Annexin V+/PI+), and necrotic (Q3; Annexin V-/PI+) populations. **(B)** Quantitative bar graphs showing the mean ± SD percentages of healthy, early apoptotic, late apoptotic, and necrotic cells across five biological replicates (n=4). Statistical comparisons between treatment groups are indicated. The results represent the mean ± SD of five independent biological replicates (n=4). Statistical significance was analyzed using two-way ANOVA followed by Tukey's post-hoc test: (ns = non-significant, * $p < 0.05$, ** $p < 0.01$, *** $p < 0.001$, **** $p < 0.0001$).

**Table SI3:** The percentage distribution of healthy cells, early apoptosis, late apoptosis, and necrosis in LoVo colon cancer cells treated with L&D@CeNPs (n=5).

| **LoVo Colon Cancer Cells** (NPs Conc. - µg/mL) | **Healty** (Mean± SD - %) | **Early Apoptosis** (Mean± SD - %) | **Late Apoptosis** (Mean± SD - %) | **Necrosis** (Mean± SD - %) |
|---|---|---|---|---|
| No treatment - 0 | 88.38 ± 5.99 | 1.58 ± 1.08 | 5.27 ± 2.57 | 4.82 ± 2.79 |
| D-Cys@CeNP - 500 | 86.65 ± 8.65 | 2.36 ± 2.15 | 4.71 ± 3.23 | 6.30 ± 3.52 |
| D-Cys@CeNP - 735 ($IC_{50}$) | 85.95 ± 8.58 | 6.83 ± 8.46 | 3.01 ± 1.93 | 4.24 ± 1.59 |
| D-Cys@CeNP - 1000 | 92.33 ± 4.10 | 1.70 ± 1.76 | 2.74 ± 2.06 | 3.22 ± 1.27 |
| L-Cys@CeNP - 500 | 84.88 ± 3.89 | 2.95 ± 2.15 | 3.85 ± 1.77 | 8.31 ± 4.67 |
| L-Cys@CeNP - 544 ($IC_{50}$) | 85.70 ± 3.03 | 4.68 ± 3.79 | 3.28 ± 1.74 | 6.33 ± 4.45 |
| L-Cys@CeNP - 1000 | 93.05 ± 5.21 | 1.36 ± 2.43 | 2.41 ± 2.37 | 3.18 ± 2.64 |

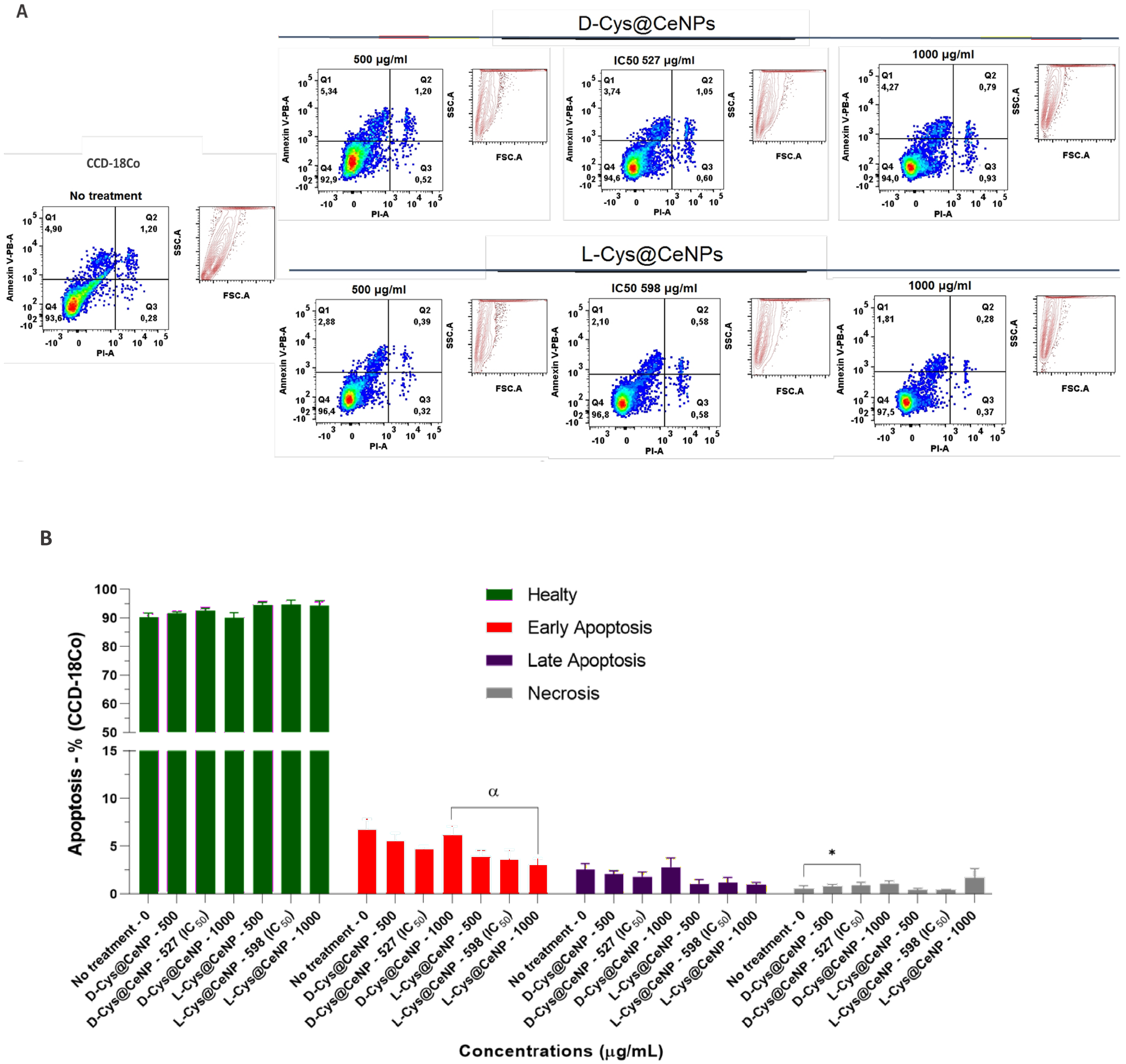

**Figure SI11. Flow cytometry-based apoptosis analysis of CCD-18Co fibroblast cells treated with D-Cys@CeNP and L-Cys@CeNP for 24 hours. (A)** Representative flow cytometry plots for each treatment condition in CCD-18Co cells. Quadrant analysis distinguishes healthy (Q4; Annexin V-/PI-), early apoptotic (Q1; Annexin V+/PI-), late apoptotic (Q2; Annexin V+/PI+), and necrotic (Q3; Annexin V-/PI+) populations. **(B)** Quantitative bar graphs showing the mean ± SD percentages of healthy, early apoptotic, late apoptotic, and necrotic cells across five biological replicates (n=4). Statistical comparisons between treatment groups are indicated. The results represent the mean ± SD of five independent biological replicates (n=4). Statistical significance was analyzed using two-way ANOVA followed by Tukey's post-hoc test: (ns = non-significant, * $p < 0.05$, ** $p < 0.01$, *** $p < 0.001$, **** $p < 0.0001$).

**Table SI 4:** The percentage distribution of healthy cells, early apoptosis, late apoptosis, and necrosis in CCD-18Co colon healthy cells treated with L&D@CeNPs (n=4).

| **CCD-18Co Colon Cancer Cells** (NPs Conc. - µg/mL) | **Healty** (Mean± SD - %) | **Early Apoptosis** (Mean± SD - %) | **Late Apoptosis** (Mean± SD - %) | **Necrosis** (Mean± SD - %) |
|---|---|---|---|---|
| **No treatment - 0** | 90.18 ± 2.96 | 6.74 ± 2.18 | 2.51 ± 1.34 | 0.59 ± 0.55 |
| **D-Cys@CeNP - 500** | 91.65 ± 0.86 | 5.51 ± 1.73 | 2.07 ± 0.70 | 0.77 ± 0.50 |
| **D-Cys@CeNP - 527 ($IC_{50}$)** | 92.58 ± 1.54 | 4.68 ± 0.62 | 1.81 ± 1.02 | 0.92 ± 0.62 |
| **D-Cys@CeNP - 1000** | 89.93 ± 3.31 | 6.16 ± 1.66 | 2.81 ± 1.64 | 1.12 ± 0.39 |
| **L-Cys@CeNP - 500** | 94.58 ± 1.84 | 3.95 ± 1.18 | 1.06 ± 0.87 | 0.43 ± 0.28 |
| **L-Cys@CeNP - 598 ($IC_{50}$ )** | 94.70 ± 3.03 | 3.63 ± 2.00 | 1.23 ± 1.01 | 0.44 ± 0.05 |
| **L-Cys@CeNP - 1000** | 94.30 ± 3.32 | 3.07 ± 1.27 | 0.98 ± 0.46 | 1.65 ± 1.96 |

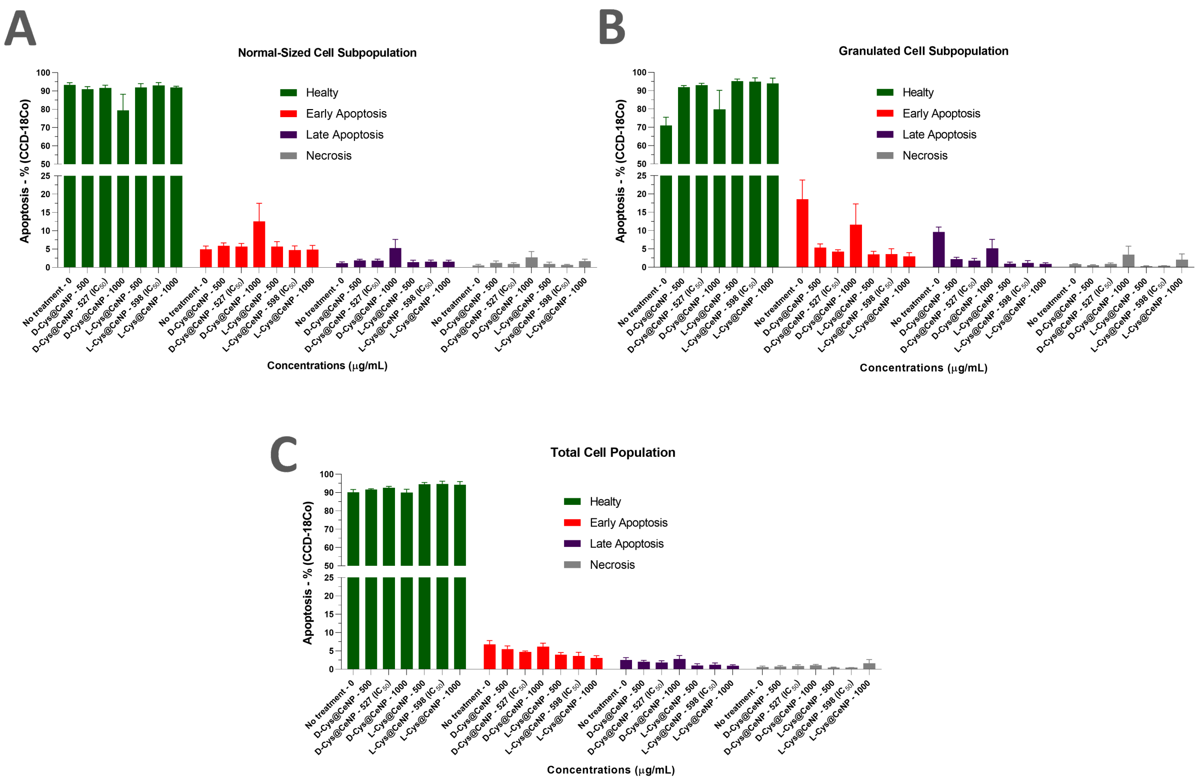

**Figure SI 12. Apoptosis analysis of granulated and normal cell populations in CCD-18Co healthy fibroblast cells.** Flow cytometry-based apoptosis assessment was performed in CCD-18Co cells, distinguishing two main subpopulations: normal cells (A) and granulated cells (B) based on FSC (Forward Scatter) and SSC (Side Scatter) parameters. Additionally, the overall apoptotic response in the total cell population (C) was evaluated. Apoptotic profiles were analyzed using Annexin V/PI staining, distinguishing healthy (Annexin V-/PI-), early apoptotic (Annexin V+/PI-), late apoptotic (Annexin V+/PI+), and necrotic (Annexin V-/PI+) populations. Panel descriptions: **(A)** Normal cells: Apoptotic distribution in cells with lower SSC values. **(B)** Granulated cells: Apoptotic distribution in cells exhibiting high SSC values. **(C)** Total cell population: Overall apoptotic response, including both granulated and normal cell populations. Data represent mean ± SD from four independent biological replicates

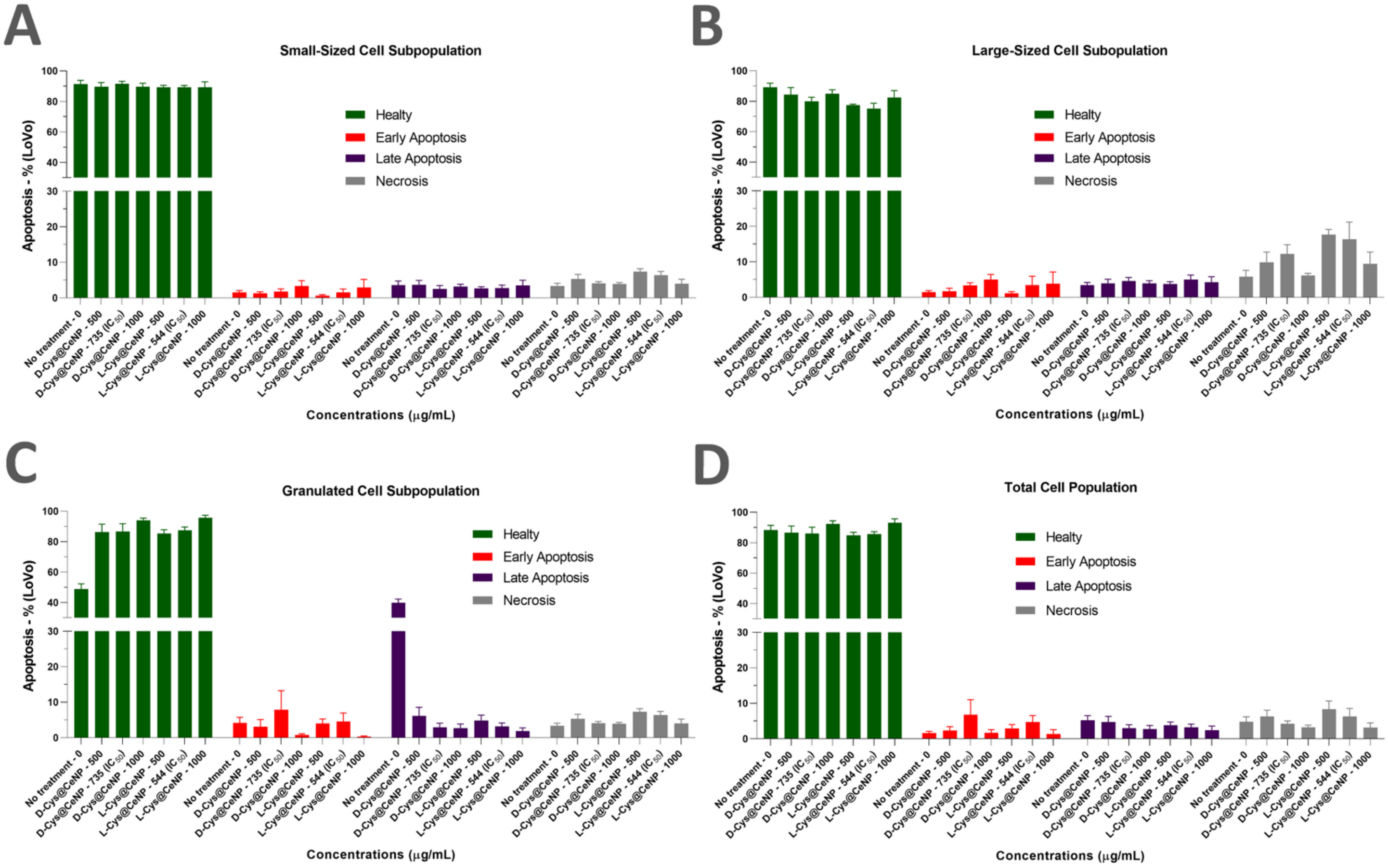

**Figure SI 13. Apoptosis analysis of distinct cell subpopulations based on FSC/SSC gating in LoVo colon cancer cells.** Flow cytometry-based apoptosis assessment was performed in LoVo cells, where three distinct subpopulations were identified based on FSC (Forward Scatter) and SSC (Side Scatter) parameters: small-sized cells **(A),** large-sized cells **(B),** and granulated cells (**C).** Additionally, the overall apoptotic response in the total cell population **(D)** was analyzed. Apoptotic profiles were evaluated using Annexin V/PI staining, distinguishing healthy (Annexin V-/PI-), early apoptotic (Annexin V+/PI-), late apoptotic (Annexin V+/PI+), and necrotic (Annexin V-/PI+) populations. Panel descriptions: **(A)** Small-sized cell population: Apoptotic distribution in cells with low FSC and SSC values. **(B)** Large-sized cell population: Apoptotic distribution in cells with high FSC values. **(C)** Granulated cell population: Apoptotic distribution in cells with high SSC values. **(D)** Total cell population: Overall apoptotic response including all subpopulations. Data represent mean ± SD from four independent biological replicates

**TNFR signaling pathway**

The canonical TNFR signaling pathway **(Figure SI14)** is a sophisticated molecular network that balances cellular survival and programmed death. The efficiency of this pathway is governed by three critical proteins, A20, NEMO, and IκBα, which together form a functional regulatory axis that determines the duration, intensity, and ultimate outcome of NF-κB activation.

**1. TNFAIP3 (A20): The Ubiquitin-Editing Brake**

A20 is a potent negative regulator and a dual-function ubiquitin-editing enzyme. It acts as the primary "brake" of the pathway by terminating the signaling cascade. Specifically, A20 dismantles K63-linked polyubiquitin chains on RIPK1 and NEMO, effectively destabilizing the signaling platform required for downstream activation. This deubiquitinase activity is essential for preventing chronic inflammation and ensuring that the NF-κB survival response is transient and tightly controlled.

**2. IKBKG (NEMO): The Essential Structural Scaffold**

NEMO (Inhibitor of Nuclear Factor Kappa-B Kinase Regulatory Subunit Gamma) serves as the indispensable structural bridge within the IKK complex. It acts as a scaffold that links upstream polyubiquitination events to the activation of catalytic kinases. Without NEMO, the IKK complex cannot assemble or phosphorylate its targets, effectively halting the transmission of the survival signal. Its presence is the prerequisite for moving the signal from the receptor to the cytoplasm.

**3. NFKBIA (IκBα): The Cytoplasmic Gatekeeper**

IκBα is the final checkpoint of the NF-κB pathway. In its stable state, it sequesters the p50/RELA heterodimer in the cytoplasm, preventing nuclear translocation. The pathway only proceeds when IκBα is phosphorylated and degraded, allowing NF-κB to enter the nucleus and drive gene transcription. IκBα also functions in a rapid feedback loop; as a target gene of NF-κB, its newly synthesized proteins quickly re-enter the cytoplasm to re-capture NF-κB, ensuring the signal is properly terminated.

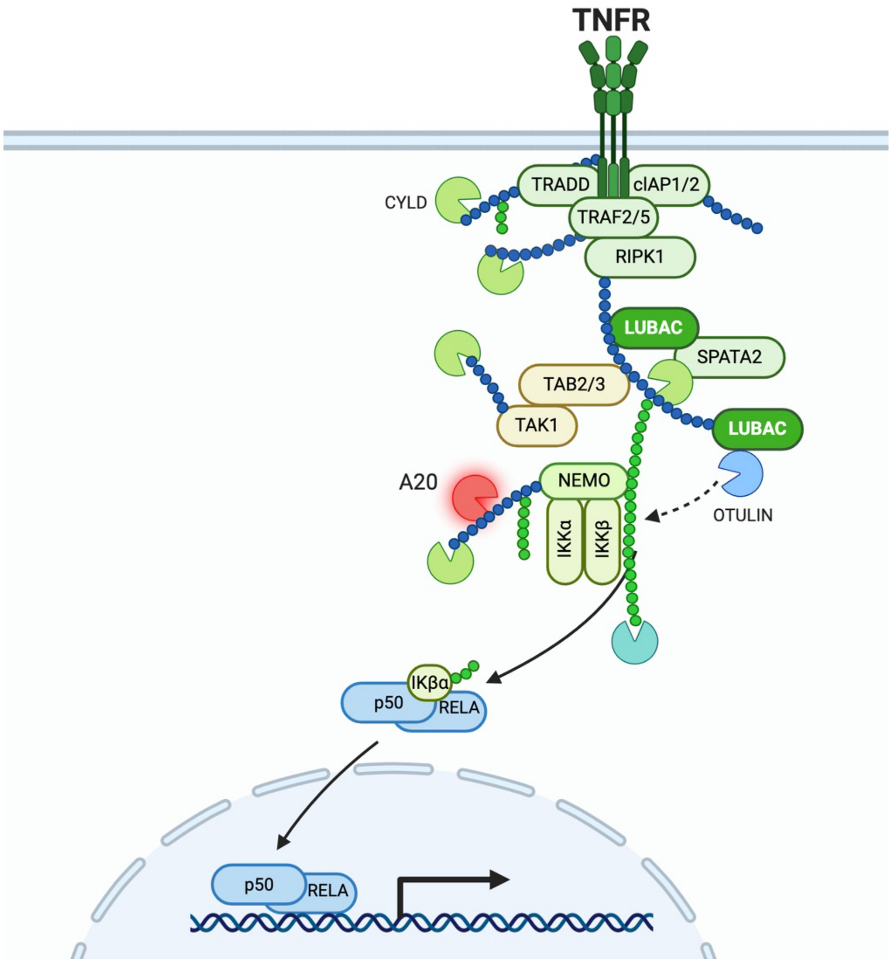

**Figure SI 14.** Signaling pathway of TNFAIP3/A20 gene and its correlation with p50 subunit of NFKB1A gene and IKB (red labeled). *Created by Biorender*.

**Table SI 5: ROS & Sensitivity Summary Table to create radar plot for composite ROS-toxicity index**

| Cell Line | D-Cys@CeNP IC50 | L-Cys@CeNP IC50 | D-Cys@CeNP ROS | L-Cys@CeNP ROS | D-Cys@CeNP Sensitivity | L-Cys@CeNP Sensitivity | D-Cys@CeNP ROS Index | L-Cys@CeNP ROS Index |
|---|---|---|---|---|---|---|---|---|
| CCD-18Co | 527 | 598 | 1.9 | 1.2 | 0.0019 | 0.00167 | 0.00361 | 0.00201 |
| LoVo | 735 | 544 | 3.8 | 1.4 | 0.00136 | 0.00184 | 0.00517 | 0.00257 |
| DLD-1 | 129 | 116 | 2.8 | 1.8 | 0.00775 | 0.00862 | 0.02171 | 0.01552 |
| COLO-201 | 30 | 60 | 3.2 | 4.2 | 0.03333 | 0.01667 | 0.10667 | 0.07 |

**SI Table 6: The values of flow cytometry populations for healthy, early and late apoptosis and necrosis columns at IC50 values for 24h.**

| CCD-18Co Colon Cancer (NPs Conc. - µg/mL) | Healty (Mean± SD - %) | Early Apoptosis (Mean± SD - %) | Late Apoptosis (Mean± SD - %) | Necrosis (Mean± SD - %) |
|---|---|---|---|---|
| D-Cys@CeNP - 527 (IC50) | 92.58 ± 1.54 | 4.68 ± 0.62 | 1.81 ± 1.02 | 0.92 ± 0.62 |
| L-Cys@CeNP - 598 (IC50 ) | 94.70 ± 3.03 | 3.63 ± 2.00 | 1.23 ± 1.01 | 0.44 ± 0.05 |
| **LoVo Colon Cancer Cells (NPs Conc. - µg/mL)** | **Healty (Mean± SD - %)** | **Early Apoptosis (Mean± SD - %)** | **Late Apoptosis (Mean± SD - %)** | **Necrosis (Mean± SD - %)** |
| D-Cys@CeNP - 735 (IC50) | 85.95 ± 8.58 | 6.83 ± 8.46 | 3.01 ± 1.93 | 4.24 ± 1.59 |
| L-Cys@CeNP - 544 (IC50) | 85.70 ± 3.03 | 4.68 ± 3.79 | 3.28 ± 1.74 | 6.33 ± 4.45 |
| **DLD-1 Colon Cancer Cells (NPs Conc. - µg/mL)** | **Healty (Mean± SD - %)** | **Early Apoptosis (Mean± SD - %)** | **Late Apoptosis (Mean± SD - %)** | **Necrosis (Mean± SD - %)** |
| D-Cys@CeNP - 129 (IC50) | 78.62 ± 9.02 | 15.66 ± 7.87 | 3.91 ± 2.21 | 1.09 ± 0.47 |
| L-Cys@CeNP - 116 (IC50) | 78.32 ± 9.56 | 15.31 ± 10.78 | 4.07 ± 1.30 | 1.50 ± 0.90 |
| **COLO-201 Colon Cancer (NPs Conc. - µg/mL)** | **Healty (Mean± SD - %)** | **Early Apoptosis (Mean± SD - %)** | **Late Apoptosis (Mean± SD - %)** | **Necrosis (Mean± SD - %)** |
| D-Cys@CeNP - 30 (IC50) | 44.44 ± 4.58 | 1.04 ± 0.44 | 16.82 ± 8.54 | 37.68 ± 10.04 |
| L-Cys@CeNP - 60 (IC50) | 36.60 ± 12.52 | 3.94 ± 4.76 | 25.88 ± 16.34 | 33.60 ± 15.94 |

**SI Table S7:** The values used to create radar graph in Figure 6; normalized table where each value in the apoptosis, necrosis, and sensitivity columns is scaled relative to the highest value in that column (i.e., maximum = 1).

| Cell Line | Nanoparticle Treatment | Early Apoptosis (%) | Late Apoptosis (%) | IC50 (µg/mL) | Necrosis (%) | Sensitivity (1/IC50) |
|---|---|---|---|---|---|---|
| **CCD-18Co** | **D-Cys@CeNP** | **0.29885057** | **0.06993818** | **527** | **0.02441614** | **0.05705706** |
| **CCD-18Co** | **L-Cys@CeNP** | **0.23180077** | **0.04752705** | **598** | **0.01167728** | **0.05105105** |
| **LoVo** | **D-Cys@CeNP** | **0.43614304** | **0.11630603** | **735** | **0.11252654** | **0.04204204** |
| **LoVo** | **L-Cys@CeNP** | **0.29885057** | **0.12673879** | **544** | **0.16799363** | **0.05405405** |
| **DLD-1** | **D-Cys@CeNP** | **1** | **0.15108192** | **129** | **0.02892781** | **0.23423423** |
| **DLD-1** | **L-Cys@CeNP** | **0.97765006** | **0.1572643** | **116** | **0.03980892** | **0.25825826** |
| **COLO-201** | **D-Cys@CeNP** | **0.06641124** | **0.64992272** | **30** | **1** | **1** |
| **COLO-201** | **L-Cys@CeNP** | **0.25159642** | **1** | **60** | **0.89171975** | **0.5015015** |

**Table SI 8:** Gene expression table to create the radar graph in Figure 7

| | | | |
|---|---|---|---|
| **CCD-18Co Colon Cancer Cells** | **TNFAIP3/A20** | **IKBKG** | **NFKBIA/P50** |
| **(NPs Conc. - µg/mL)** | **Fold Chance** | **Fold Chance** | **Fold Chance** |
| **D-Cys@CeNP - 527 (IC50)** | **0.76** | **1.89** | **1.43** |
| **L-Cys@CeNP - 598 (IC50 )** | **1.04** | **2.17** | **2.68** |
| **LoVo Colon Cancer Cells** | **TNFAIP3/A20** | **IKBKG** | **NFKBIA/P50** |
| **(NPs Conc. - µg/mL)** | **Fold Chance** | **Fold Chance** | **Fold Chance** |
| **D-Cys@CeNP - 735 (IC50)** | **1.61** | **3.51** | **3.33** |
| **L-Cys@CeNP - 544 (IC50)** | **2.15** | **3.01** | **2.03** |
| **DLD-1 Colon Cancer Cells** | **TNFAIP3/A20** | **IKBKG** | **NFKBIA/P50** |
| **(NPs Conc. - µg/mL)** | **Fold Chance** | **Fold Chance** | **Fold Chance** |
| **D-Cys@CeNP - 129 (IC50)** | **0.33** | **1.93** | **0.64** |
| **L-Cys@CeNP - 116 (IC50)** | **0.61** | **3.18** | **0.55** |
| **COLO-201 Colon Cancer Cells** | **TNFAIP3/A20** | **IKBKG** | **NFKBIA/P50** |
| **(NPs Conc. - µg/mL)** | **Fold Chance** | **Fold Chance** | **Fold Chance** |
| **D-Cys@CeNP - 30 (IC50)** | **2.6** | **0.75** | **1.3** |
| **L-Cys@CeNP - 60 (IC50)** | **0.75** | **2.12** | **2.84** |

**Gene Expression Profiling via qPCR**

To investigate the potential anticancer mechanisms of D-Cys@CeNPs and L-Cys@CeNPs, gene expression profiling was performed using quantitative PCR (qPCR) to analyze the expression levels of TNFAIP3, IKBKG, and NFKBIA, with β-actin serving as the housekeeping control gene. RNA isolation and cDNA synthesis were performed prior to qPCR analysis, and the primers designed for these genes are listed in Table 1.

**Table SI 9.** Forward and Reverse Primer seguences of TNFAIP3, IKBKG and NFKBIA

| Gene of Interest | Forward and Reverse Primers | | Product Length(bp) |
|---|---|---|---|
| **TNFAIP3 (A20)** | Forward<br>Reverse | GGGGCCCGGAGAGGTAAC<br>CCAGTGTGTATCGGTGCATGG | 197 |
| **IKBKG (NEMO** | Forward<br>Reverse | CCCCTCACTCCCTGTGAAGC<br>TACGTCCTGATCTGCTGCCG | 140 |
| **NFKBIA (IκBα)** | Forward<br>Reverse | GGAAGTGATCCGCCAGGTGA<br>CTCCCAGAAGTGCCTCAGCA | 125 |

**Total RNA Extraction and cDNA Synthesis:**

For cDNA synthesis, 1 µg of total RNA was used with the iScript cDNA Synthesis Kit (BioRad Laboratories, CA, USA, Cat# 1708891). The reaction components and volumes were prepared according to the kit protocol, and the synthesis steps are detailed in Tables 2 and 3. The total reaction volume, including oligo(dT) and random primers, was set to 20 µL, and cDNA synthesis was carried out in a thermal cycler following the protocol outlined in these tables.

**Table SI10.** Reaction components and volumes in cDNA synthesis.

| **Component** | **Volume per Reaction (µL)** |
|---|---|
| 5× iScript Reaction Mix | 4 |
| iScript Reverse Transcriptase | 1 |
| Nuclease-free water | Variable |
| RNA template (100 ng total RNA) * | Variable |
| **Total** | **20** |

*A maximum of 1µg of RNA was used for a total reaction volume of 20 µL.

**Table SI11.** Reaction protocols for Reverse Transcription.

| Steps | Temperature(°C) | Time (minute) |
|---|---|---|
| Priming | 25 | 5 |
| Reverse Transcription | 46 | 20 |
| RT inactivation | 95 | 1 |
| Optional step | 4 | - |

**RT-qPCR Analysis:**

cDNA samples (10 ng/µL) were amplified using the SYBR Green PCR Master Mix (Bio-Rad, Cat# 1725120). Real-time PCR was performed on the AriaMx Real-Time PCR System (Agilent, USA) under standard cycling conditions. Gene expression levels were quantified relative to β-actin as the housekeeping gene using the $2^{-\Delta\Delta Ct}$ method. The kit components and qPCR steps are detailed in Tables 4 and 5.

**Table SI12.** Reaction compound for qPCR.

| Reagent | Final Concentration | Amount (µL) |
|---|---|---|
| SYBR Green PCR Master Mix | | 10 |
| Forward Primer | 5 µM | 1 |
| Reverse Primer | 5 µM | 1 |
| cDNA Template | 40 ng | 4 |
| $dH_2O$ | | 4 |
| **Total Reaction Volume** | | **20** |

**Table SI13.** Amplification curves for qPCR analysis.

| Step | Temperature (°C) | Duration | # of Cycles |
|---|---|---|---|
| DNA Denaturation | 95 | 2 min | 1 |
| Denaturation | 95 | 10 sec | 40 |
| Annealing/Extension | 58 | 30 sec | 40 |
| Extension | 72 | 30 sec | 40 |
| Final Extension | 72 | 10 min | 1 |
| Hold | 4 | - | - |
